# Organization of the corticotropin-releasing hormone and corticotropin-releasing hormone-binding protein systems in the central nervous system of the sea lamprey *Petromyzon marinus*

**DOI:** 10.1101/2022.04.22.489136

**Authors:** Daniel Sobrido-Cameán, Laura González-Llera, Ramón Anadón, Antón Barreiro-Iglesias

## Abstract

The expression of the corticotropin-releasing hormone (*PmCRH*) and the CRH-binding protein (*PmCRHBP*) mRNAs was studied by *in situ* hybridization in the brain of prolarvae, larvae and adults of the sea lamprey *Petromyzon marinus*. We also generated an antibody against the PmCRH mature peptide to study the distribution of PmCRH-immunoreactive cells and fibers. PmCRH immunohistochemistry was combined with anti-tyrosine hydroxylase immunohistochemistry, *PmCRHBP in situ* hybridization or neurobiotin transport from the spinal cord. The most numerous PmCRH-expressing cells were observed in the magnocellular preoptic nucleus-paraventricular nucleus and in the superior and medial rhombencephalic reticular formation. *PmCRH* expression was more extended in adults than in larvae, and some cell populations were mainly (olfactory bulb) or only (striatum, ventral hypothalamus, prethalamus) observed in adults. The preopto-paraventricular fibers form conspicuous tracts coursing towards the neurohypophysis, but many immunoreactive fibers were also observed coursing in many other brain regions. Brain descending fibers in the spinal cord come from cells located in the isthmus and in the medial rhombencephalic reticular nucleus. The distribution of *PmCRHBP*-expressing neurons was different from that of PmCRH cells, with cells mainly present in the septum, striatum, preoptic region, tuberal hypothalamus, pretectum, pineal complex, isthmus, reticular formation, and spinal cord. Again, expression in adults was more extended than in larvae. *PmCRH*- and *PmCRHBP*-expressing cells are different, excluding colocalization of these substances in the same neuron. Present findings reveal a complex CRH/CRHBP system in the brain of the oldest extant vertebrate group, the agnathans, which shows similarities but important divergences with that of mammals.

## 1. Introduction

Corticotropin-releasing hormone (CRH) [also known as corticotropin-releasing factor (CRF)], is a neuropeptide consisting of 41 amino acids that was first purified from sheep hypothalamus (Vale et al., 1981). CRH is involved in stress response as the major regulator of the hypothalamo-pituitary-adrenal stress axis (Ketchesin et al., 2017a; Yuan et al., 2019). In non-mammalian vertebrates, CRH also has a potent thyrotrophic function (see De Groef et al., 2006). In addition to its role as a neurohormone, CRH has widespread roles within the brain as a neuropeptide involved in various functions (for example, memory enhancement: Todorovic et al., 2005; anxiogenic and antinociceptive effects: Miguel and Nunes-de-Souza, 2011; stress control: Bale et al., 2000, Fox and Lowry, 2013; Yuan et al., 2019). CRH actions at synapses are mediated by CRH receptors and the CRH-binding protein (CRHBP), which are necessary for the transduction of the perception of stressful environmental situations into survival-based physiology and behavior (Behan et al., 1996; Jaferi and Bhatnagar, 2007; Ketchesin et al., 2017a).

The CRH system possibly appeared early in metazoan evolution because homologue CRH-like peptides, CRH receptors and the CRHBP have been found both in invertebrates and vertebrates (see Lovejoy and Lannoy, 2013; Cai et al., 2021). In vertebrates, the CRH-family genes are widely distributed along the nervous system in all studied species (Cardoso et al., 2016, 2020). Initial studies in the sea lamprey *Petromyzon marinus* revealed the existence of a CRH system with three CRH family members [named as CRH, urotensin I (UI) and urocortin 3 (Ucn3)], two types of CRH receptors (named alpha and beta) and a single CRHBP (Endsin, 2013; Endsin et al., 2017). These authors showed, by using polymerase chain reaction (PCR) methods, that all the components of a functional CRH system are expressed in the sea lamprey brain, and these genes are regulated in response to stress, suggesting that a hypothalamo-pituitary-interrenal axis is functional in lampreys. More recently, Cardoso and coworkers (2020) reported the presence of five CRH-family members in the sea lamprey [named CRH/UCNa (corresponds to CRH of Endsin et al., 2017), CRH/UCNb (corresponds to UI of Endsin et al. 2017), CRH/UCNc, UCNa (corresponds to Ucn3 of Endsin et al., 2017 and Sobrido-Cameán et al., 2021a), and UCNb]. Although Cardoso et al. (2020) could not infer clear orthology relationships with all gnathostome members of the CRH family in their phylogenetic and synteny analyses, CRH of Endsin et al. (2017)/CRH/UCNa of Cardoso et al. (2020) corresponds to the gnathostome CRH1 (Endsin et al. 2017; Cardoso et al., 2020). The pattern of expression of Ucn3 (UCNa of Cardoso et al., 2020) in the sea lamprey brain was recently reported by means of *in situ* hybridization (ISH) by our group (Sobrido-Cameán et al., 2021a), but the brain/spinal cord distribution of the other members of the CRH system is not known.

Lampreys are interesting as animal model for comparative studies due to the position that they occupy in the vertebrate phylogeny, the lamprey lineage together with myxines (hagfishes) belongs to the agnatha, the sister out-group of gnathostomes (Forey and Janvier, 1993; Delarbre et al., 2002; Furlong and Holland, 2002; Murakami et al., 2005; Kuratani and Ota, 2008). The brain of lampreys exhibits the basic cell types, regions, and structures of the vertebrate brain (Murakami and Kuratani, 2008; Lammana et al., 2022). Research on the nervous, endocrine, and immune systems of lampreys has significance for revealing the origin and evolution of these systems, to infer the ancestral organization of the brain in vertebrates and could contribute to a better understanding of human diseases and treatments (see Barreiro-Iglesias and Rodicio, 2012; Sobrido-Cameán and Barreiro-Iglesias, 2022). Here, we studied by ISH the expression of the *Petromyzon marinus* CRH (Endsin et al., 2017; in the following named PmCRH) and CRHBP (Endsin et al., 2017; PmCRHBP) mRNAs in the brain/spinal cord of sea lampreys at different life stages, as well as the distribution of neurons and fibers expressing the mature PmCRH peptide by immunohistochemistry, to better understand the development and organization of the CRH system in a jawless vertebrate.

## 2. Material and methods

### 2.1. Animals

Prolarvae (n = 15), larvae (n = 20), downstream migrating young adults (post-metamorphic juveniles; n = 6) and upstream migrating adults (n = 8) of sea lampreys, *Petromyzon marinus* L., were used for this study (see Supplementary Table 1 for details of the animals used in each experiment). Prolarvae were obtained from *in vitro* fertilized eggs reared in our laboratory (their ages are indicated as days posthatching). Downstream migrating juveniles and larvae (ammocoete; lengths comprised between 80 and 110 mm., 4-7 years old) were collected from the River Ulla (Galicia, Spain) with permission from the *Xunta de Galicia*. Upstream migrating adults were acquired from local suppliers. Adults/juveniles were fixed immediately upon arrival to the laboratory, and larvae were maintained in aquaria containing river sediment and with appropriate feeding, aeration, and temperature conditions until the day of use. Before all experiments, animals were deeply anesthetized with 0.1% tricaine methanesulfonate (MS-222; Sigma-Aldrich, St. Louis, MO, USA) in fresh water and killed by decapitation. All experiments were approved by the Bioethics Committee at the University of Santiago de Compostela and the Xunta de Galicia (project reference 15012/2020/011) and were performed in accordance with European Union and Spanish guidelines on animal care and experimentation.

### 2.2. Cloning and sequencing of the PmCRH and PmCRHBP precursor cDNAs

Larvae were anesthetized by immersion in MS-222 (Sigma; see above) and the brain and spinal cord were dissected out under sterile conditions. Total RNA was isolated from these tissues using the TriPure reagent (Roche, Mannhein, Germany). The first-strand cDNA synthesis reaction from total RNA was catalyzed with Superscript III reverse transcriptase (Invitrogen, Waltham, MA, USA) using random primers (hexamers; Invitrogen). For PCR cloning, specific oligonucleotide primers (forward: 5’-CCACCAGCCTTCTCGTCCTC-3’; reverse: 5’- CCGATGCTCGTTCTCCTCGT-3’) were designed based on the *PmCRH* precursor cDNA sequence that is deposited in GenBank with accession number KX446863.1 (Endsin et al., 2017); and specific oligonucleotide primers (forward: 5’-GCTGGAGCTGTTGACGATGC-3’; reverse: 5’-CTCGGGGCTAGAAGGGAACG-3’) were designed based on the putative *PmCRHBP* cDNA sequence that is deposited in GenBank with accession number KX446865.1 (Endsin et al., 2017). The amplified fragments were cloned into pGEM-T easy vectors (Promega, Madison, WI, USA) using standard protocols and sequenced by GATC Biotech (Cologne, Germany). Sequencing confirmed that we cloned fragments (*PmCRH*: 305 bp; *PmCRHBP*: 341 bp) of the same sequences deposited in GenBank by Endsin et al. (2017).

### 2.3. In situ hybridization

Templates for *in vitro* transcription were prepared by PCR amplification as follows. A fragment of the *PmCRH* and *PmCRHBP* precursor sequence was obtained using the primers mentioned above. But in this case, the reverse primer included the sequence of the universal T7 promoter (TAAGCTTTAATACGACTCACTATAGGGAGA). For the generation of sense probes, the sequence of the T7 promoter was included in the forward primers. Digoxigenin (DIG)-labeled riboprobes were synthesized using the amplified fragments as templates and following standard protocols using a T7 polymerase (Nzytech, Lisbon, Portugal).

ISH experiments were performed as previously described for other neuropeptide riboprobes (Sobrido-Cameán et al., 2019, 2020b, 2021a, 2021c). Briefly, whole prolarvae or the brains/rostral spinal cords of larvae and young and mature adults were dissected out and fixed by immersion for 12 hours in 4% paraformaldehyde (PFA) in phosphate-buffered saline (PBS) at 4 °C. Then, they were cryoprotected with 30% sucrose in PBS, embedded in Tissue Tek (Sakura, Torrance, CA, USA), frozen in liquid nitrogen-cooled isopentane, and cut serially on a cryostat (14 µm thickness) in transverse planes. Sections were mounted on Superfrost® Plus glass slides (Menzel, Braunschweig, Germany). The sections were incubated with DIG-labeled antisense riboprobes (1 µg/mL) at 70 °C overnight in hybridization mix and treated with RNase A (Sigma) in post-hybridization washes. Then the sections were incubated with a sheep anti-DIG antibody conjugated to alkaline phosphatase (1:2000; Roche) overnight at 4 °C. Staining was conducted in BM Purple (Roche) at 37 °C until the signal was clearly visible. Finally, the sections were mounted in Mowiol® (Sigma). No staining was observed when incubating brain sections with sense probes.

### 2.4. Generation of a novel anti-PmCRH antibody

A modified mature PmCRH peptide (CSDEPPISLDLTFHLLREVLEMAKAEQLAQQAHTNRQIMENI-NH2) with the addition of a cysteine residue to the N-terminus was synthesized by Biomedal (Sevilla, Spain) to enable coupling to key limpet hemocyanin (KLH). A rabbit (8 weeks old, New Zealand White) was first immunized subcutaneously with the peptide-KLH conjugate (400 mg) emulsified in Freund’s complete adjuvant. After four weeks, the rabbit was immunized once weekly for 2 weeks by intramuscular injection of the peptide-KLH conjugate (200 mg) emulsified in Freund’s complete adjuvant. Pre-immune (negative control with no anti-PmCRH antibodies present) and post-immunization bleeds were collected. 2.5 mL of antiserum from the final bleed were purified using a protein A-Sepharose column (GE Healthcare, Little Chalfont, UK) and the purified antibodies were used for the immunohistochemical analyses.

### 2.5. Tissue processing for immunohistochemistry

For immunohistochemistry, the whole prolarvae or brains/spinal cords of larvae and juveniles and adults were fixed by immersion in 4% PFA in 0.05 M Tris-buffered saline pH 7.4 (TBS) for 4 to 12 hours at 4 °C. The samples were then rinsed in TBS, cryoprotected with 30% sucrose in TBS, embedded in Tissue Tek (Sakura), frozen in liquid nitrogen-cooled isopentane, and cut serially on a cryostat (14-20 µm thickness) in transverse planes. Sections were mounted on Superfrost® Plus glass slides (Menzel).

### 2.6. PmCRH and TH double immunofluorescence experiments

Some samples were incubated with the purified rabbit polyclonal anti-PmCRH antibody (dilution 1:1500) at 4 °C for 72 hours. Other samples were processed for double immunofluorescence experiments using a cocktail of a mouse monoclonal anti-tyrosine hydroxylase (TH) antibody (dilution 1:1000; Millipore, Temecula, CA; Cat# MAB318; lot 0509010596; RRID: AB_2201528; immunogen: TH purified from PC12 cells) in combination with the anti-PmCRH antibody. Primary antibodies were diluted in TBS containing 15% normal goat serum and 0.2% Triton as detergent.

For double detection of PmCRH and TH by indirect immunofluorescence, the sections were rinsed in TBS and incubated for 1 hour at room temperature with a cocktail of Cy3-conjugated goat anti-rabbit (1:200; Millipore; Cat# AP132C; RRID:AB_92489) and FITC-conjugated goat anti-mouse (1:100; Millipore; Cat# AQ303F; RRID:AB_92818) antibodies. Sections were rinsed in TBS and distilled water and mounted with Mowiol® (Sigma).

### 2.7. Double in situ hybridization and immunohistochemistry

In some samples, ISH for *PmCRHBP* was followed by immunohistochemistry against PmCRH. These experiments were performed as described in a previous study (Sobrido-Cameán et al., 2020a). Briefly, after the ISH signal was clearly visible, sections were rinsed twice in TBS and treated with 10% H_2_O_2_ in TBS for 30 minutes. For heat-induced epitope retrieval, sections were treated with 0.01 M citrate buffer (pH 6.0) for 30 minutes at 90°C and allowed to cool for 20 minutes at RT. Then, the sections were incubated the anti-PmCRH primary antibody solution (same as above) overnight at RT. After rinsing in TBS, the sections were incubated with the solution containing the secondary antibody goat anti-rabbit IgG serum HRP conjugated (Dako, Glostrup, Denmark; Cat# P0448, RRID: AB_2617138, dilution 1:200) for 1 hour at room temperature. The immunoreaction was developed with 0.25 mg/ml diaminobenzidine tetrahydrochloride (DAB; Sigma) containing 0.00075% H_2_O_2_. Finally, the sections were mounted in Mowiol® (Sigma).

### 2.8. Antibody characterization

The anti-serum and the purified fraction of anti-PmCRH antibody were tested for specificity with an enzyme-linked immunosorbent assay (ELISA) using standard methods. The antiserum, the purified antibody (at concentrations from 1:1000 to 1:100000) and the pre-immune serum (1:1000) were tested against the synthetic PmCRH peptide without KLH (at a concentration of 1 mM) in the ELISA. Both the antiserum and the purified fraction gave a positive signal at all the tested concentrations, while the very low signal registered with the pre-immune serum was the same as the one obtained in the negative controls with PBS only. Since the pattern of PmCRH-immunoreactive (-ir) neuronal populations in the central nervous system (CNS) matches the pattern of *PmCRH* transcript expressing populations seen by ISH (see results), it further supports the specificity of the antibody for this peptide.

The specificity of the anti-TH antibody was tested by the supplier. The anti-TH antibody was also tested in Western blots of sea lamprey and rat brain protein extracts in our laboratory, which revealed bands of similar size for the sea lamprey and rat TH enzymes (Barreiro-Iglesias et al., 2008b). This antibody has been used in many immunohistochemical studies of lamprey brain and retina (Pierre et al., 1997; Villar-Cerviño et al., 2006; Barreiro-Iglesias et al., 2010a, 2017).

As a general control for the secondary antibodies, some sections were processed as above, except that the primary antiserum was omitted. No staining was observed in these controls.

### 2.9. Retrograde tract-tracing study of spinal cord-projecting PmCRH-ir neurons

The origin of PmCRH-ir fibers observed in the spinal cord was investigated by neuronal tract tracing combined with immunofluorescence. Tract-tracing experiments were performed in larval samples to label descending neurons that innervate the spinal cord. Neurobiotin (MW 322.8 Da; Vector; Burlingame, CA) was used as a tracer. The larval spinal cord was exposed by a longitudinal incision made in the dorsal region of the body at the level of the fifth gill and completely cut with Castroviejo scissors. The tracer was applied in the rostral stump of the transected spinal cord with the aid of a minute pin (#000). The animals were allowed to recover at 19.5 °C with appropriate aeration conditions for 7 days to allow transport of the tracer from the application point to the neuronal soma of descending neurons. Brains of these larvae were fixed with 4% PFA and processed for PmCRH immunofluorescence as above. After the immunofluorescence protocol, the sections were incubated at room temperature with Avidin D-FITC conjugated (dilution 1:1000; Vector; Cat#: A-2001) diluted in TBS containing 0.3% Triton X-100 for 4 hours to reveal the presence of neurobiotin. Slides were rinsed in TBS and distilled water and mounted with Mowiol® (Sigma).

### 2.10. Nuclear counterstain

Nuclear counterstain was carried out after the immunofluorescence procedure by immersing the slides in 0.5 µg/mL bisbenzimide (Sigma) in TBS for 10 seconds before mounting.

### 2.11. Image acquisition and montage

Confocal photomicrographs were taken with a TCS-SP2 spectral confocal laser microscope (Leica Microsystems, Wetzlar, Germany). Data was acquired by use of appropriate laser lines and narrow spectral windows tuned to the specific absorption and emission wavelengths of each fluorescent marker (bisbenzimide, FITC or Cy3). Confocal projections of stacks were done with the LAF suite (Leica) or with ImageJ (Schneider et al., 2012). Fluorescence and bright-field photomicrographs were obtained with an AX70 epifluorescence microscope equipped with a DP70 digital camera (Olympus, Tokyo, Japan). Plates of photomicrographs and minimal bright/contrast adjustments were done with Photoshop 2021 (Adobe, San Jose, CA). Schematic drawings were done with CorelDraw 2019 (Corel, Ottawa, Canada).

## 3. Results

### 3.1. Distribution of CRHergic cell populations in the brain of sea lampreys

We studied the distribution of CRHergic cell populations in the brain and spinal cord of the sea lamprey using specific anti-*PmCRH* riboprobes for ISH and antibodies against the mature PmCRH peptide for immunofluorescence. Both ISH and immunohistochemistry revealed similar PmCRH-expressing populations distributed along the brain. The most conspicuous PmCRH-expressing populations were observed in the preoptic region and isthmus, whereas in other brain regions positive cells were much more restricted. Combination of ISH or immunochemistry for PmCRH and TH immunohistochemistry was also used to better define preoptic and hypothalamic PmCRH positive populations. With immunohistochemistry, PmCRH-ir) fibers were observed in most brain regions although at different densities, extending also in the spinal cord. The cells of origin of spinal PmCRH-ir brain descending fibers were studied by combining tract-tracing from the spinal cord (at the level of the 5^th^ gill opening) with PmCRH immunohistochemistry.

Schematic drawings of transverse sections of brains of adults and larvae based on *PmCRH* ISH results are presented in Figures 1 and 2. Photomicrographs of sections of adult brains processed by *PmCRH* ISH and PmCRH immunohistochemistry are presented in Figures 3-4 and Figure 5, respectively. Photomicrographs of sections of larval brains processed for *PmCRH* ISH and PmCRH immunohistochemistry are presented in Figure 6 and Figure 7, respectively. The distribution of PmCRH-expressing cells projecting to the spinal cord is presented in the Figure 8. Figure 9 shows photomicrographs of sections of prolarval heads processed for *PmCRH* ISH, *PmCRHBP* ISH or for PmCRH immunohistochemistry. The terminology employed in this study for the various regions, nuclei, tracts, and identified neurons followed those used in other studies of lamprey neuropeptides of our group (see Sobrido-Cameán et al., 2019, 2020b, 2021a). Since the same positive populations found by *PmCRH* ISH were also observed by PmCRH immunofluorescence, we assumed that the antibody recognized only this PmCRH peptide. We named the cells showing the *PmCRH* mRNA and the PmCRH peptide as PmCRH-expressing cells. However, the possibility that the anti-PmCRH antibody may cross-react with other related members of the CRH-family peptides (for example, CRH/UCNb of Cardoso et al., 2020) cannot be ruled out. As expected, immunohistochemistry provided additional information on the cell shape and revealed the pattern of innervation of the hypophysis and brain by PmCRH-ir fibers.

**Figure 1.**
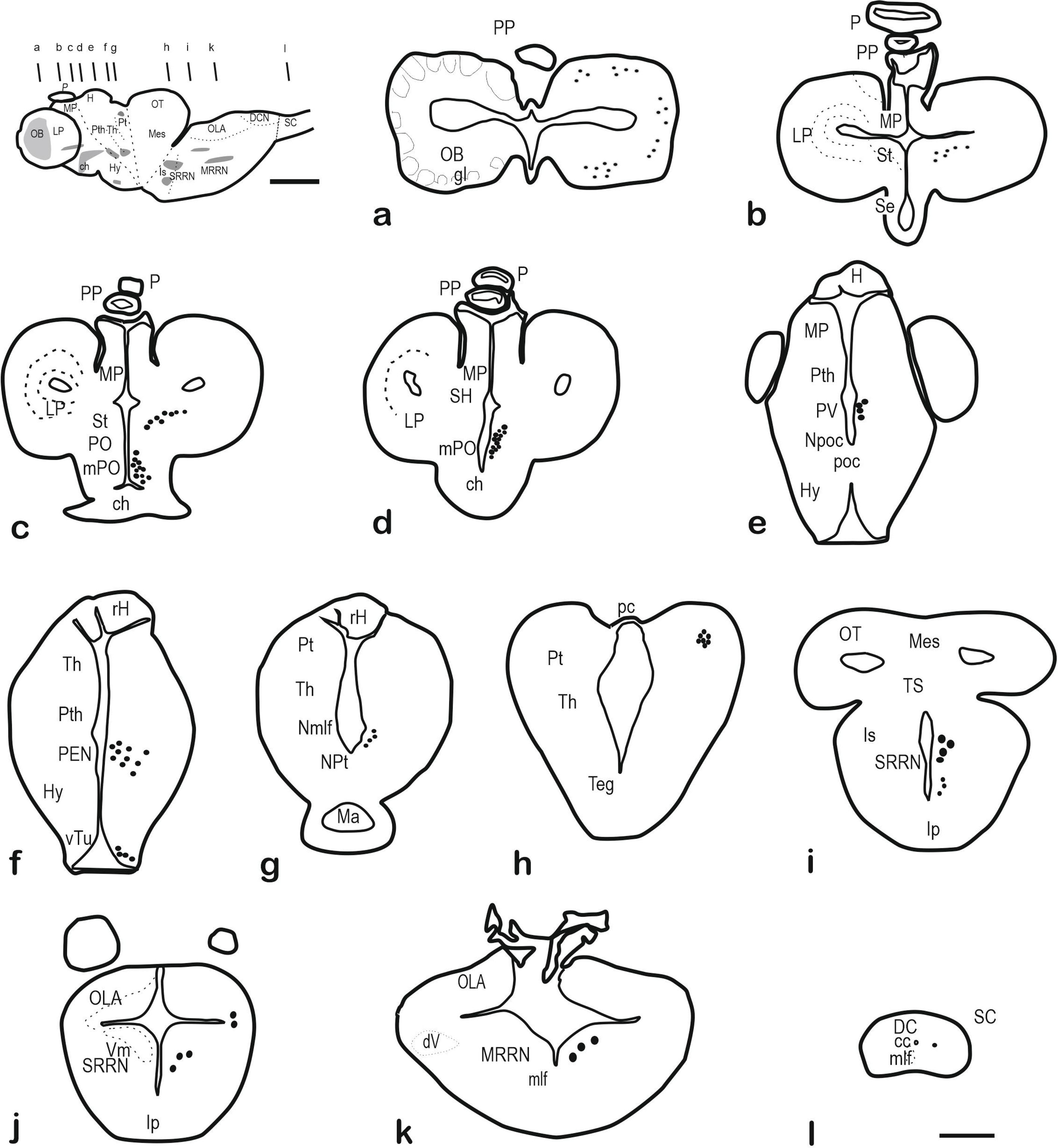
Schematic drawings of transverse sections of the adult sea lamprey brain showing the distribution of *PmCRH*-expressing neurons revealed by ISH (at the right) and the anatomical references (at the left). The levels of sections are indicated in the figurine of the lateral view of the adult brain. For abbreviations, see the list. Scale bar, 200 µm (a-l), 1 mm (figurine).

**Figure 2.**
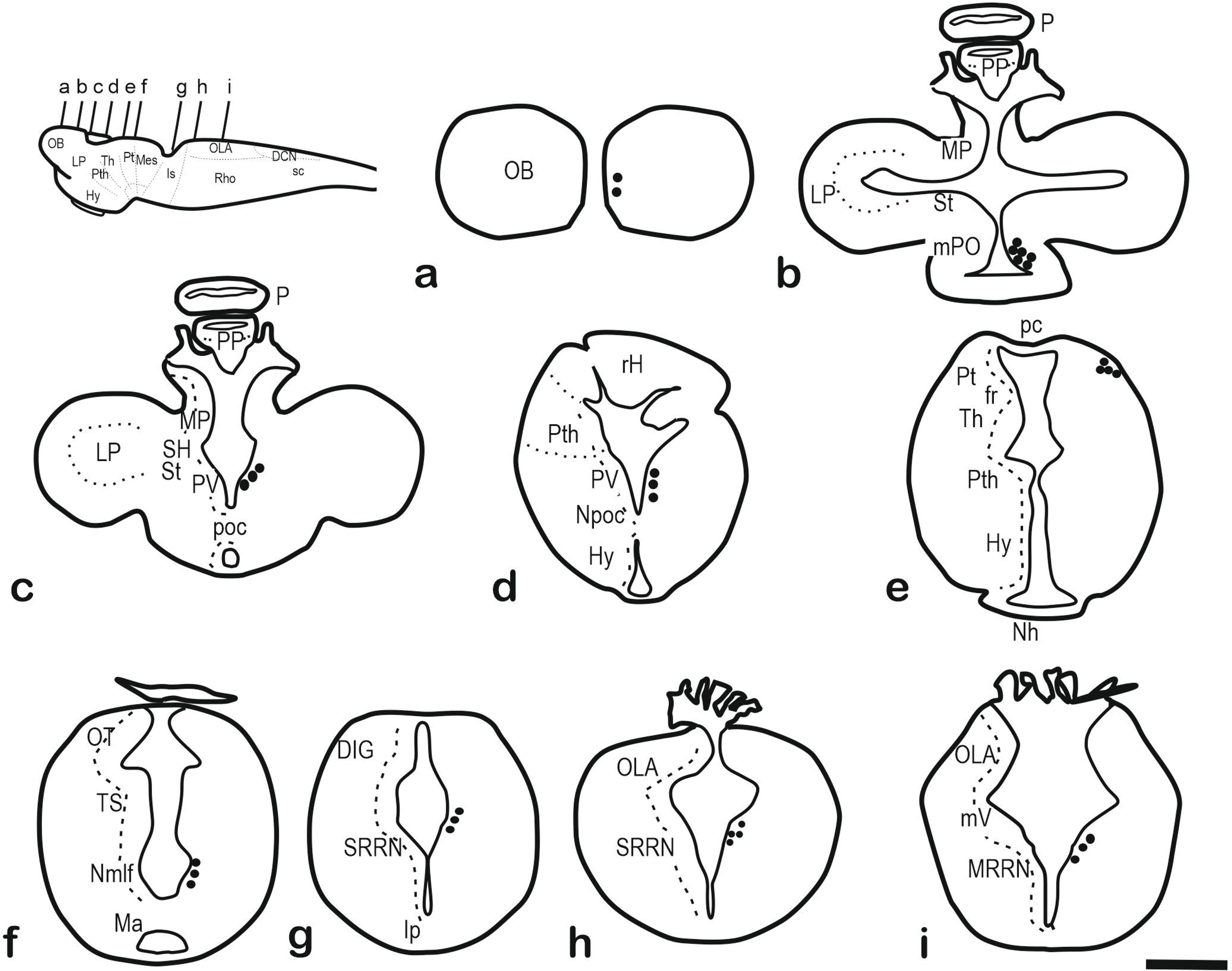
Schematic drawings of transverse sections of the brain of a larval sea lamprey showing the distribution of *PmCRH*-expressing neurons revealed by ISH (at the right) and the anatomical references (at the left). The levels of sections are indicated in the figurines of the lateral view of the brains. For abbreviations, see the list. Scale bar, 200 µm.

**Figure 3.**
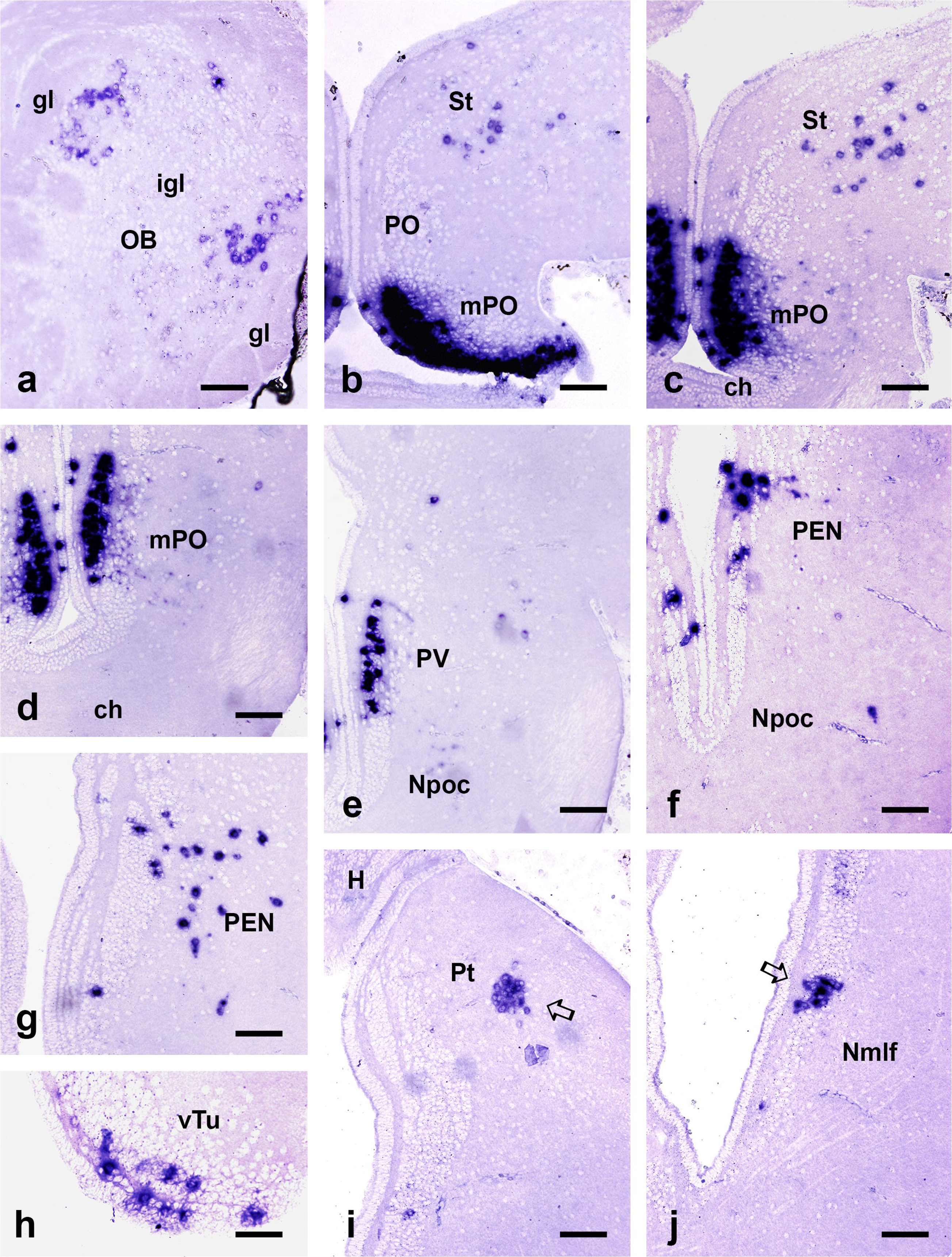
Photomicrographs of transverse sections of the forebrain of a young sea lamprey showing the expression of *PmCRH* mRNA in neurons by ISH. **a**, Groups of PmCRH-expressing neurons in the olfactory bulb located near the glomerular layer. **b-d**, Sections through the ventral telencephalon and preoptic region showing high expression in neurons of the mPO nucleus and positive neurons scattered in the striatum. **e**, Section showing positive neurons in the paraventricular nucleus, which is continuous with the mPO. **f-g**, Sections showing positive neurons in the posterior entopeduncular nucleus. **h**, Section showing positive neurons in the ventral portion of the tuberal nucleus rostral to the neurohypophysis. **i**, Section through the rostral pretectum showing a compact group of positive cells located in a central region (outlined arrow). **j**, Section through the caudal diencephalon showing a group of positive neurons in the nucleus of the medial longitudinal fascicle (outlined arrow). For abbreviations, see the list. Scale bars, 50 µm.

**Figure 4.**
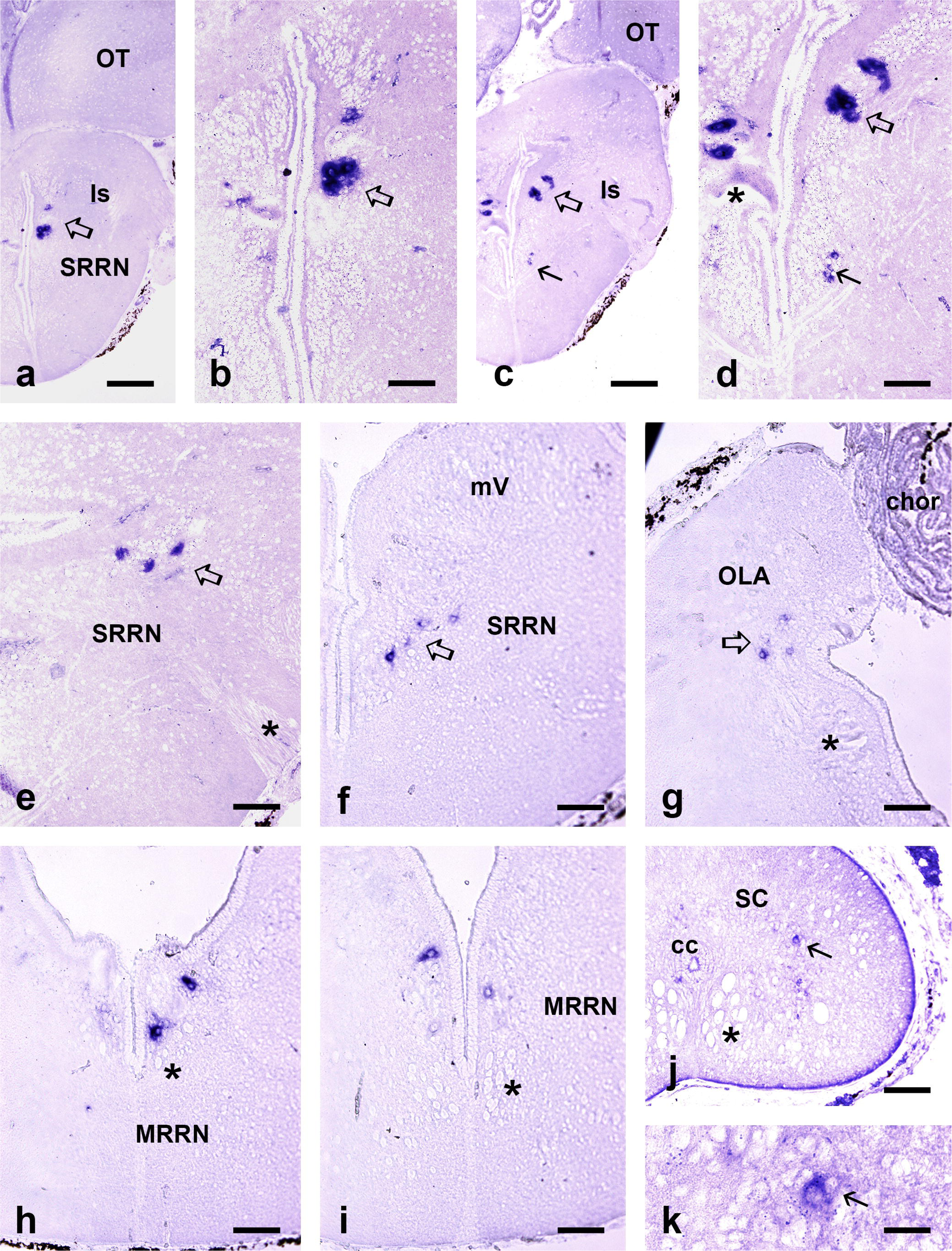
Photomicrographs of transverse sections of the hindbrain and spinal cord of a young sea lamprey showing the expression of *PmCRH* mRNA in neurons by ISH. **a-d**, Sections through the isthmus and details showing groups of large PmCRH-expressing neurons in the dorsal region (outlined arrows) and small neurons in the ventral region (arrows in c-d). The asterisk in d indicates a small portion of the giant isthmic neuron I1. **e-f**, Sections at the level of entrance of the trigeminal nerve (asterisk in e) and the trigeminal motor nucleus (mV in f) showing positive neurons (outlined arrows) in lateral and medial reticular regions, respectively. **g**, Section showing positive neurons (outlined arrow) in the octavolateralis region at the level of the glossopharyngeal motor nucleus (asterisk). **h-i**, Sections showing positive neurons in the medial rhombencephalic reticular nucleus at rostral and caudal levels, respectively. Asterisks, giant axons of the medial longitudinal fascicle. **j-k**, Section and detail showing a positive neuron (arrows) in the rostral spinal cord. Asterisk in j, medial longitudinal fascicle. For abbreviations, see the list. Scale bars, 50 µm (b, d-j), 125 µm (a, c), 12.5 µm (k).

**Figure 5.**
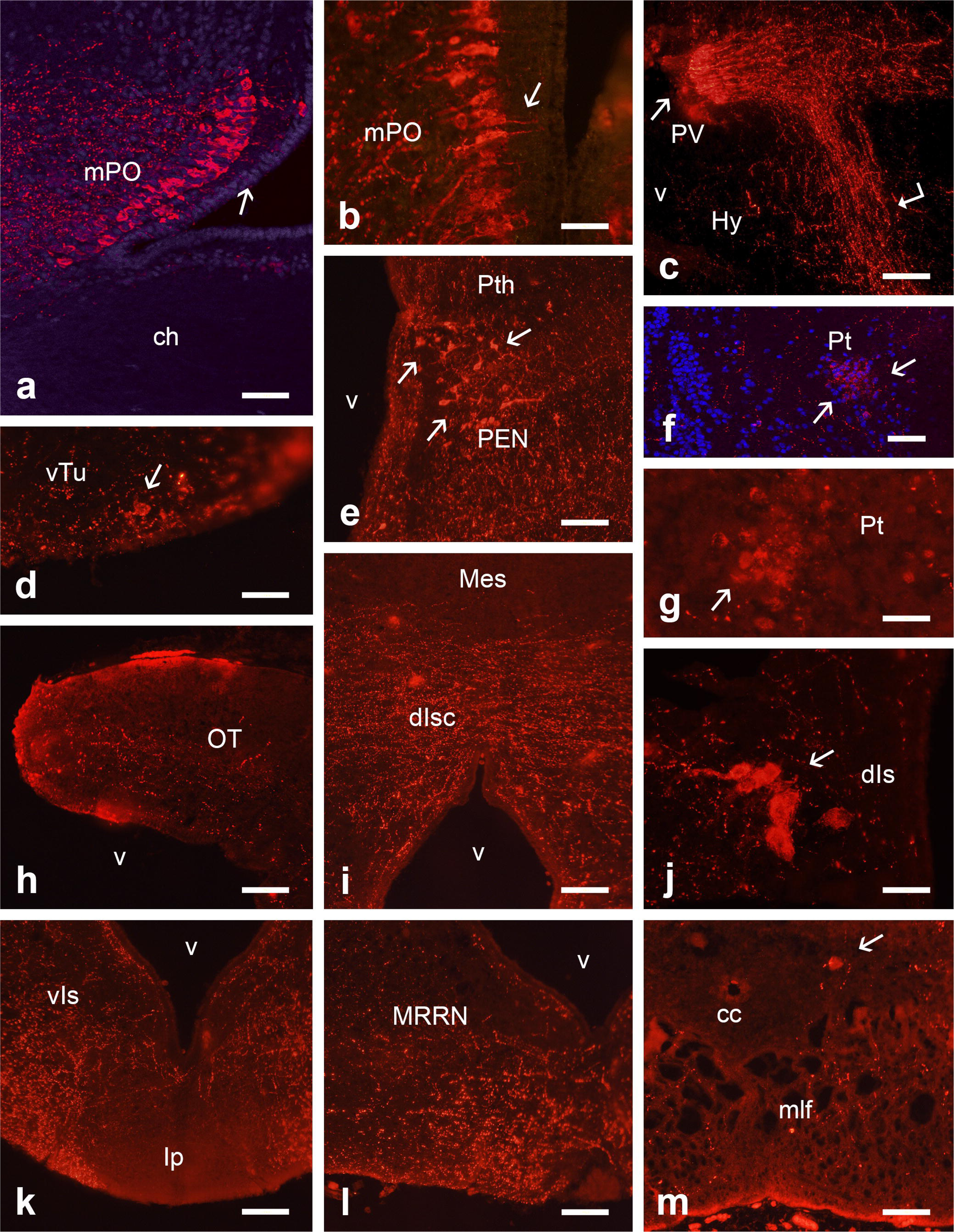
Fluorescent photomicrographs of selected sections of the brain of an adult sea lamprey showing PmCRH-ir cells and fibers. **a-b**, Photomicrograph, and detail, of the mPO nucleus. Arrow points to medial dendrites. **c**, Section showing neurons in the paraventricular nucleus (arrow) and the projection towards the hypophysis (angled arrow). **d,** Section through the ventral tuberal region showing small positive neurons (arrow). **e**, Section showing a scattered group of neurons in the posterior entopeduncular nucleus/prethalamus region (arrows) and numerous fibers. **f-g**, Sections through the pretectum showing a compact group of cells (arrows) in the central region. **h**, Section through the optic tectum showing PmCRH-ir fibers in its inner half. **i**, Section through the dorsal limit between the midbrain and hindbrain showing numerous positive fibers in the dorsal isthmic commissure. **j**, Section though the group of large positive neurons (arrow) in the dorsal isthmus near the giant isthmic cell. **k**, Section through the ventral isthmus showing abundant positive fibers except in the interpeduncular nucleus neuropil. **l**, Section of the hindbrain tegmentum showing many positive fibers near the midline caudal to the interpeduncular nucleus. **m**, Section of the rostral spinal cord showing a positive neuron (arrow) and low density of positive fibers. a and f are confocal photomicrographs with a blue nuclear contrast. In a, d, and j medial is at the right, in c, e, f, and h medial is at the left. For abbreviations, see the list. Scale bars, 50 µm (a, c, d, e, f, h, i, j, k, l, m), 25 µm (b, g).

**Figure 6.**
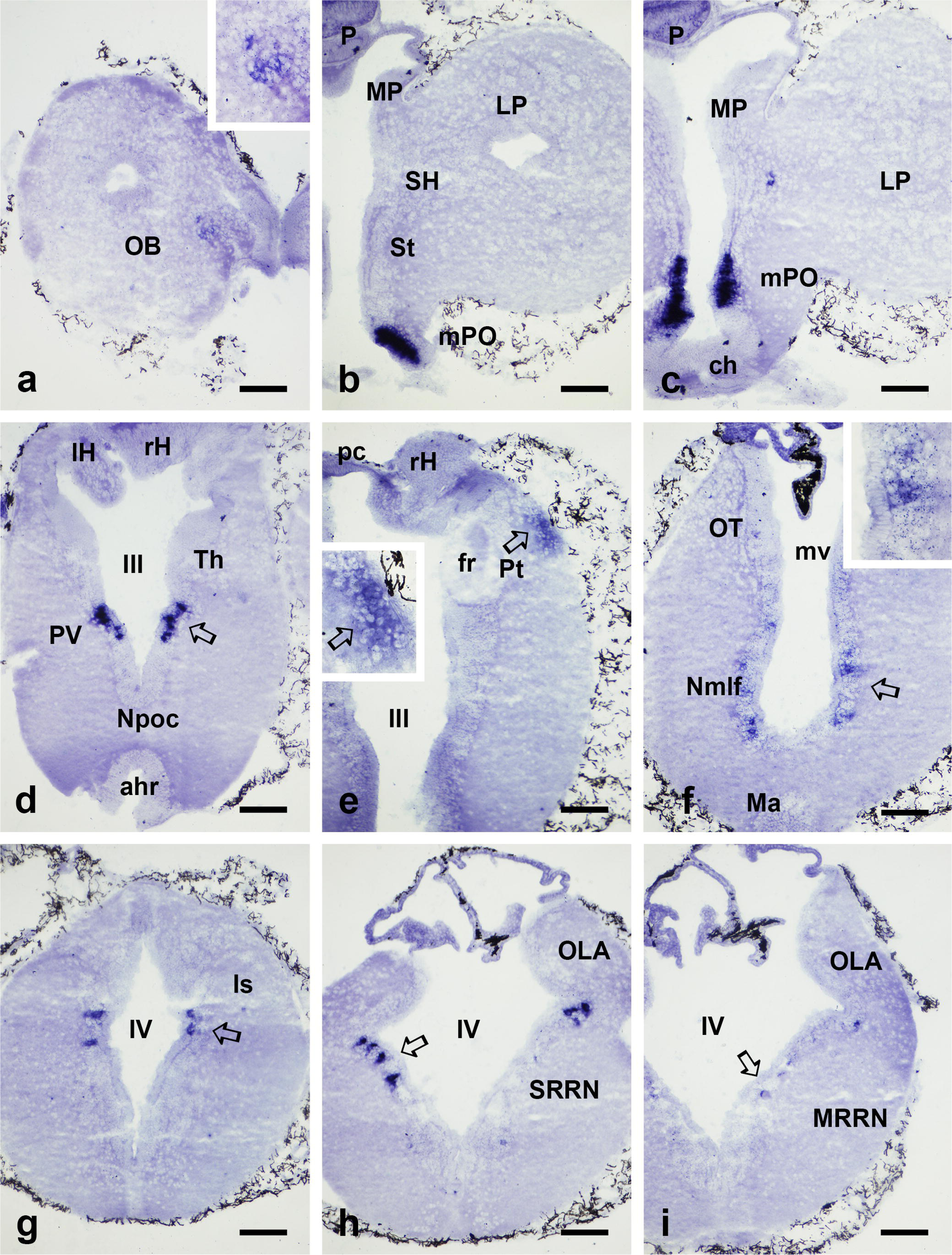
Photomicrographs of transverse sections of the brain of a lamprey larva (100 mm long) showing the distribution of *PmCRH*-expressing neurons by ISH. **a**, Section of the left olfactory bulb (OB) showing some positive neurons in the medial glomerular region. Inset, detail of cells. **b-c**, Sections of the forebrain showing positive neurons in the mPO nucleus. **d**, Section showing positive neurons in the paraventricular nucleus (outlined arrow). **e**, Section at the level of rostral pretectum showing a group of positive neurons near the meningeal surface (outlined arrow). Inset, detail of these pretectal cells. **f**, Section at the level of the nucleus of the medial longitudinal fascicle showing positive neuron (outlined arrow). Inset, detail of these cells. **g-h**, Sections of the isthmus at rostral (g) and caudal (h) levels showing positive cells in the superior rhombencephalic reticular nucleus (outlined arrows). **i**, Section at the level of the facial motor nucleus showing scarce positive neurons (outlined arrow) in the medial rhombencephalic reticular nucleus. For abbreviations, see the list. Scale bars, 50 µm.

**Figure 7.**
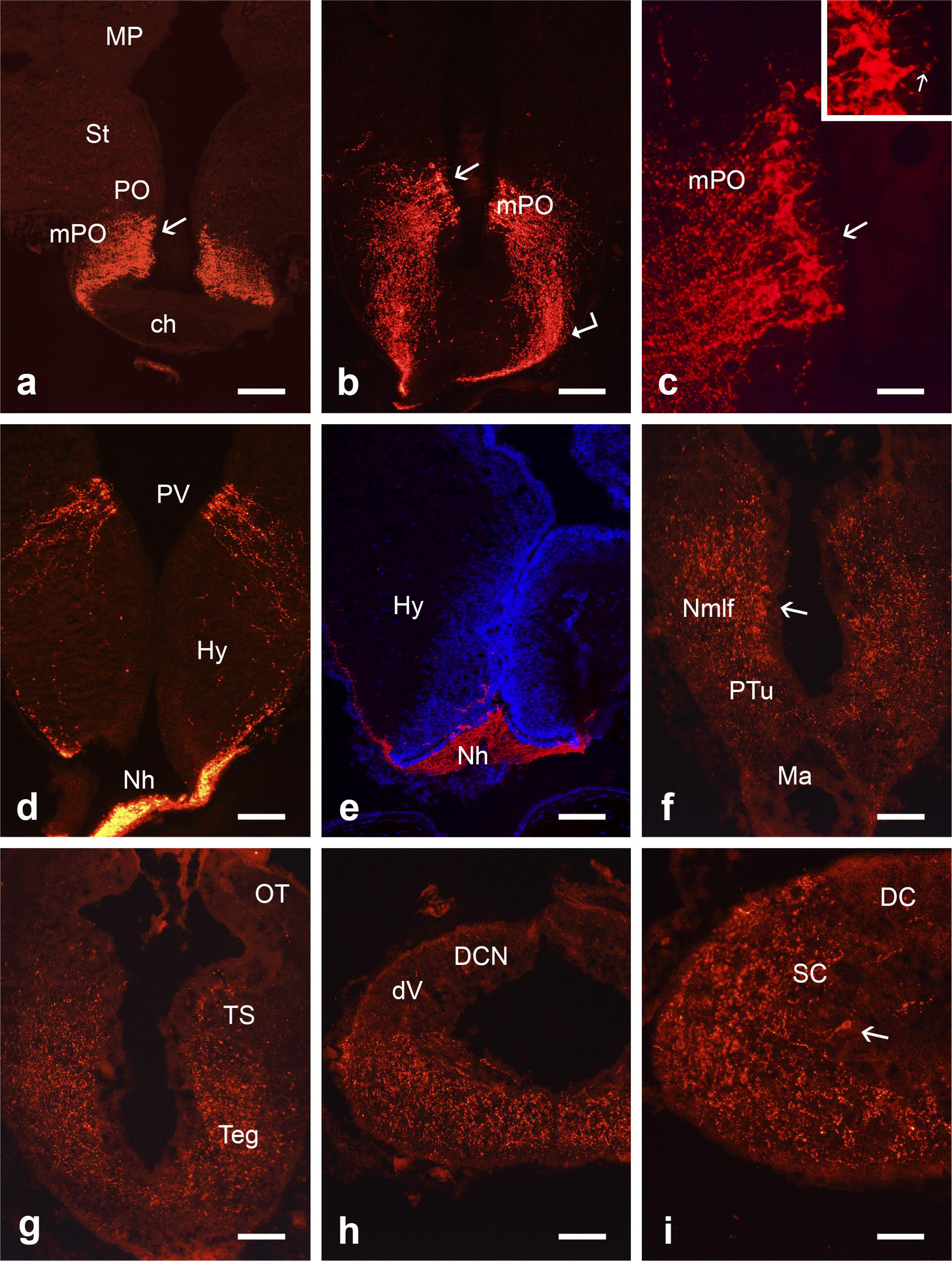
Fluorescence photomicrographs of transverse sections of the larval brain showing the distribution of cells and fibers immunoreactive for PmCRH. **a-c**, Sections of the mPO nucleus at rostral and a more caudal level showing cell bodies (arrows) and the fiber tract (angled arrow) coursing toward the neurohypophysis. In inset, note medial processes (arrow) of positive neurons. **d**, Section through the paraventricular nucleus and rostral neurohypophysis. **e**, Confocal photomicrograph of a section through the hypothalamus showing accumulation of immunoreactive fibers in the posterior neurohypophysis. Note the scant fibers in the hypothalamus. In blue, nuclear contrast. **f**, Section through the diencephalon showing immunoreactive cells (arrow) in the nucleus of the medial longitudinal fascicle and many positive fibers. **g,** Section through the midbrain showing abundant PmCRH-ir fibers in the tegmentum and torus semicircularis. **h**, Section through the caudal hindbrain showing abundant fibers in basal regions but scant PmCRH-ir fibers in the alar region (dV and DCN). **i**, Section through the rostral spinal cord showing abundant PmCRH-ir fibers and a PmCRH-ir neuron (arrow). For abbreviations, see the list. Scale bars, 50 µm (a, b, d, e, f, g, h), 25 µm (c, i).

**Figure 8.**
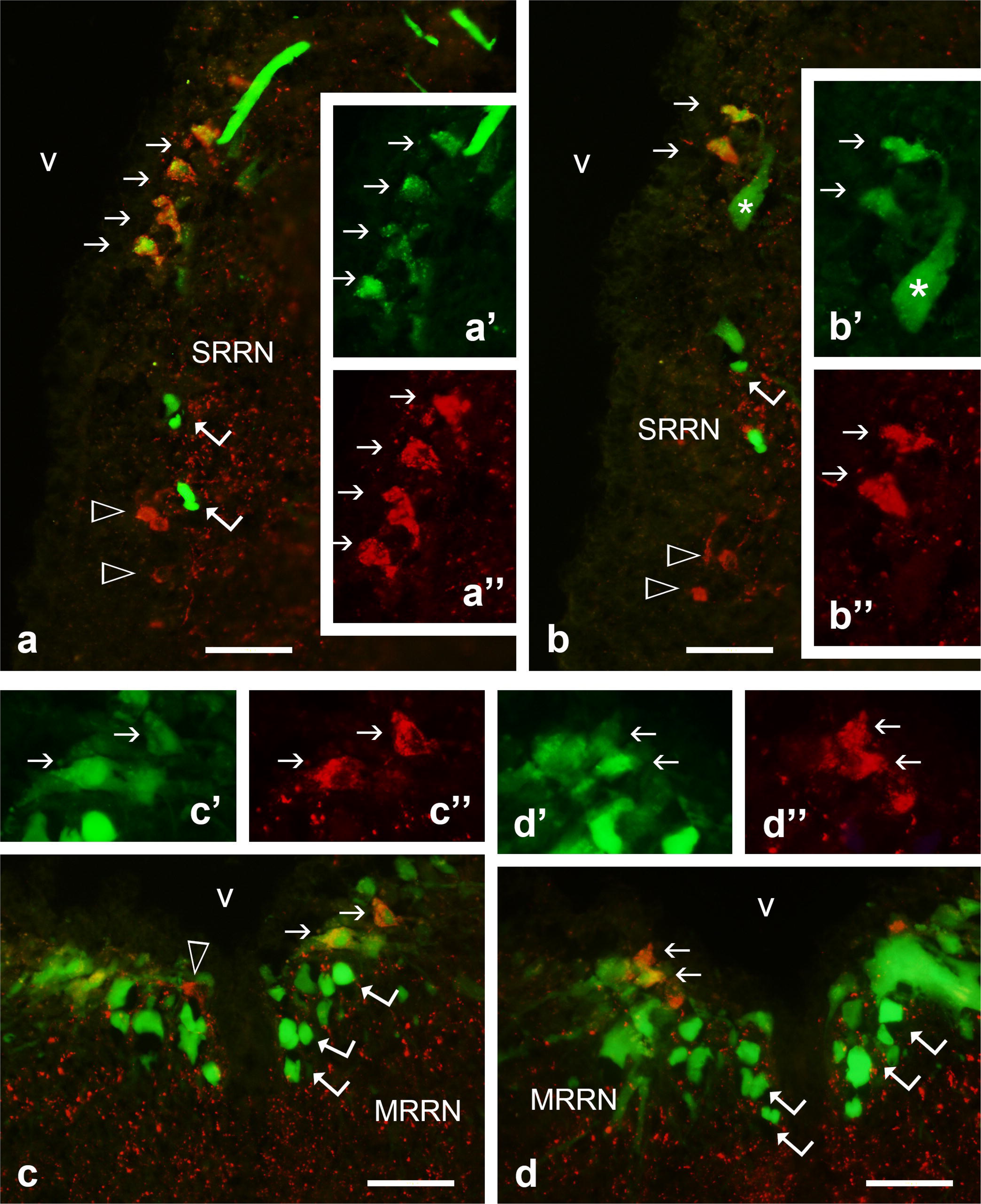
Photomicrographs showing PmCRH-ir neurons in the rhombencephalon retrogradely labeled after injection of neurobiotin in the spinal cord. **a-b**, Cells in the superior rhombencephalic reticular formation (isthmus) double-labeled for PmCRH and neurobiotin (arrows), axons of giant neurons labeled with neurobiotin (angled arrows), and PmCRH-ir cells (arrowheads). Insets (a’, a’’, b’ and b’’), detail of double labeled cells in single channels (red, PmCRH; green: neurobiotin). Asterisk in b-b’ indicates a dendrite of a giant isthmic neuron. **c-d**, Double-labeled neurons (arrows) in the middle rhombencephalic reticular formation among numerous perikarya and axons (angled arrows) of neurobiotin - labeled reticulospinal cells. Insets (c’, c’’, d’, d’’), single channel details of double labeled cells. Red channel, PmCRH; green channel: neurobiotin. For abbreviations, see the list.

**Figure 9.**
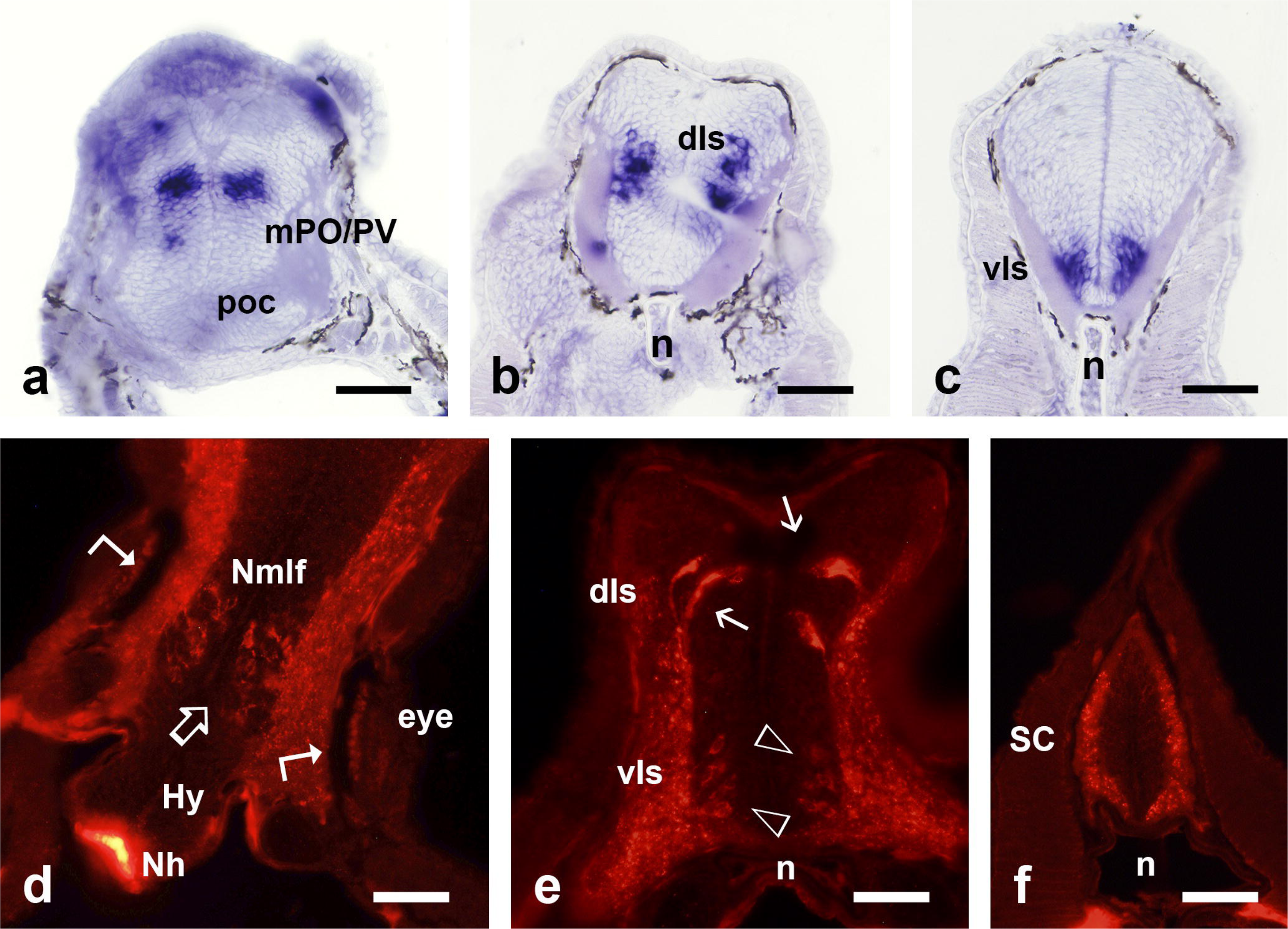
Photomicrographs of transverse sections of lamprey prolarvae showing the distribution of neurons expressing PmCRH mRNA (a-c) and PmCRH immunoreactivity (d-f). **a**, Section through the forebrain of a P25 prolarvae showing a group of PmCRH positive neurons in the primordium of the mPO/paraventricular nucleus dorsal to the postoptic commissure. **b and c**, Sections of the hindbrain of a P25 prolarvae showing groups of positive neurons in the dorsal isthmus (b) and ventral region caudal to the otic vesicle (c). Note the notochord (n) at these levels. **d**, Section of the caudal forebrain of a P30 prolarvae at the level of the eyes showing PmCRH-immunoreactive neurons in the mantle layer (Nmlf, outlined arrow) and numerous positive fibers in the marginal layer neuropil. Note also intense immunoreactivity in the primordium of the neurohypophysis (Nh). Angled arrows point to the pigment cell layer of the retina, which is close to the brain surface. **e**, Section at the level of the isthmus of a P30 prolarvae showing large PmCRH-ir dorsal neurons (arrows) and smaller and paler ventral neurons (arrowheads). Note also abundant fibers in the marginal layer. **f,** Section through the spinal cord of a P30 prolarvae showing abundant PmCRH-ir fibers in the marginal layer. For abbreviations, see the list. Scale bars, 50 µm (a, b. c), 25 µm (d, e, f).

#### 3.1.1. Adult lampreys

In the olfactory bulbs, both in postmetamorphic juveniles and upstream migrating adults, PmCRH-expressing cells are scattered or in small groups in the granular layer close the glomerular layer (Figures 1a, 3a). These cells are small or medium sized, but the specific cell type could not be determined. The topographical relation of these groups with glomeruli varied around the bulb, some glomeruli showing associated conspicuous *PmCRH*-expressing populations and other glomeruli lacking them or showing scarce positive cells.

In the telencephalic lobes of postmetamorphic juveniles and upstream migrating adults, a band of *PmCRH*-expressing cells was observed extending between the lateral part of the striatum and the ventral region of the pallium, away from the ventricle (Figures 1b-c, 3b-c). These cells do not occupy the dense band of striatal neurons but lie externally to it (for a characterization of the lamprey striatum, see Pombal et al., 1997).

The most conspicuous PmCRH-expressing population in juvenile/adult lampreys was observed in the preoptic region. A large group of PmCRH-expressing cells with high staining intensity by ISH and immunofluorescence was observed in the magnocellular preoptic nucleus (mPO) of juvenile and adult sea lampreys (Figures 1b-c, 3b-d, 5a-b), whereas the medial preoptic nucleus that separates it from the PmCRH-expressing striatal population lacks PmCRH-expressing cells. The band of PmCRH-expressing mPO cells extends from the dorsolateral (caudal) zone to the medial wall of the preoptic recess forming a dense population. Most cells are in a cell band separated by a layer of neuropil from the subependymal cells, which only shows scarce PmCRH-expressing cells. Combination with TH immunohistochemistry reveals that TH-ir neurons and PmCRH-expressing cells occupy ventral and dorsal locations in the wall of lateral preoptic recesses, respectively, without intermixing (not shown). PmCRH immunohistochemistry reveals that the labeled mPO cells send lateral processes that extend to the hypothalamus ventrocaudally towards the neurohypophysis (Figure 5a-b). In addition, these cells (or at least a part of them) send a medial process toward the surface of the preoptic recess (Figure 5a-b). The band of PmCRH-expressing mPO cells extends caudally from the dorsal region of the optic recess to the paraventricular nucleus (Figures 1e, 3e, 5c), which shows cells like those of the mPO also sending medial processes toward the ventricle and ventrolateral processes to the same lateroventral hypothalamic fiber region. The conspicuous band of PmCRH-ir axons coursed through the ventrolateral hypothalamus, which showed a large density of these fibers (Figure 5c).

In the hypothalamus of juvenile/adult lampreys, PmCRH-expressing small cells were also observed in the ventral tuberal nucleus rostral to the neurohypophysis (Figures 1f, 3h, 5d). These small cells are scattered near the ventricle.

The lamprey diencephalon is comprised of three prosomeres (p1-p3; Pombal and Puelles, 1999). In the diencephalon of juveniles and upstream migrating adults, a band of PmCRH-expressing cells were scattered in the ventral (anterior) region of the prethalamus (p3)/posterior entopeduncular nucleus (Figures 1f, 3f-g, 5e), with some cells near to or accompanying the hypothalamic paraventricular organ. This PmCRH-expressing population does not extend caudally till the conspicuous dopaminergic nucleus of the posterior tubercle (as seen with combined TH immunohistochemistry, not shown). The location of cells of this population in the hypothalamus or prethalamus is difficult to assess, since they are intermingled with most dorsal TH-ir CSF-contacting cells of the paraventricular organ (hypothalamic), but some cells are in the thick cell layers typical of the prethalamus, just lateral to the paraventricular organ, or widely scattered in the neuropil lateral to it forming a loose population. The location of these scattered cells may correspond in part to the posterior entopeduncular area or posterior hypothalamus of Pombal and Puelles (1999).

In the pretectum (alar p1) of juveniles and adults, a small compact group of PmCRH-expressing cells was observed in a dorsocentral region of neuropil and cells (Figures 1h, 3i, 5f-g). The location of this PmCRH-expressing group is like that of a group of pretectal cells that projects to the habenula and parapineal ganglion reported with tracing methods (see discussion), but the possible identity will need to be assessed experimentally. Some PmCRH-expressing small neurons were observed also in the basal region of p1 (nucleus of the medial longitudinal fascicle) (Figures 1g, 3j).

In the rhombencephalon (hindbrain) of juveniles/adults, the most conspicuous PmCRH-expressing cells were observed in the isthmus (Figures 1i, 4a-d, 5j), but sparser populations of PmCRH-expressing reticular neurons were observed along almost the entire hindbrain (Figures 1j-k, 4e-i,). In the isthmus, a group of large or medium-sized neurons strongly expressing PmCRH was observed close to the giant Müller isthmic cell (I1) (Figures 1i, 4a-d). More ventrally, some small positive cells were also observed (Figure 4c-d). The difference in size of dorsal and ventral cells was remarkable (Figures 4c-d, 5j-k). From just caudal to the trigeminal motor nucleus, a scattered population of PmCRH-expressing neurons was observed along most of the hindbrain medial to visceromotor nuclei or in the medial reticular region (Figures 1j-k, 4e-i). These PmCRH-expressing neurons were small or medium-sized. Only occasional PmCRH-expressing cells were observed in the rostral spinal cord, close to the obex (Figures 1-l, 4j-k, 5m). No PmCRH cells were observed at more caudal spinal cord levels (at the level of the 5^th^ gill, midbody region or most caudal spinal cord; Supplementary Figure 1a-b).

#### 3.1.2. Larval and prolarval lampreys

The larval telencephalon was almost devoid of PmCRH-expressing cells. In the olfactory bulbs of larvae of about 100 mm in length (3-4 years old), a few PmCRH-expressing cells were observed associated to medial glomeruli, with other bulbar regions lacking such cells (Figures 2a, 6a). No positive cells were observed in the olfactory bulb primordia of prolarvae examined, or in the striatum and pallium of larvae and prolarvae.

The mPO-paraventricular population of *PmCRH*-expressing cells is conspicuous in larvae of around 100 mm in length (Figures 2b, 6b-c, 7a-c). As in adults, many axons coursed toward the neurohypophysis, and medial dendrites directed toward the ventricle are also observed in many positive cells (Figure 7a-e). In 15-days posthatching (P15) prolarvae, the preopto-paraventricular population was not observed yet with ISH or immunofluorescence, but in P25-P30 prolarvae a group of *PmCRH*-expressing cells was observed in sections of the prosencephalon at transverse levels just dorsorostral to the postoptic commissure (Figure 9a). These sections show the ventral surface of the brain next to the preoral epithelium without intervening notochord and a scant marginal layer outside the comparatively thick mantle. PmCRH immunofluorescence in P25 and P30 prolarvae showed positive cells in this mPO/paraventricular nucleus location (not shown) and intense fluorescence in the primordial neurohypophysis (Figure 9d). By comparison of sections containing *PmCRH*-expressing cells with those of previous studies of our group with GABA, PCNA, and HNK-1 immunohistochemistry (Meléndez-Ferro et al., 2002a, 2003; Villar-Cheda et al., 2006), we identify this population as a primordium of the preopto-paraventricular population observed in older life stages. Unlike in adults, no PmCRH expression was observed in cells of the ventral hypothalamus of larvae or prolarvae.

In larvae or prolarvae, no prethalamo-PEN-hypothalamic PmCRH-expressing population was observed. In the pretectum of larvae of about 100 mm in length, a small compact group of faint stained *PmCRH*-expressing cells was observed located close to the meninges lateral to the posterior commissure organ (Figures 2e, 6e), instead of occupying a central position in the pretectum as observed in adults. Some PmCRH-expressing small neurons were also observed in the nucleus of the medial longitudinal fascicle of larvae (Figures 2f, 6f, 7f) and in P30 prolarvae (Figure 9d). No PmCRH-expressing cell populations were observed in the thalamus or the midbrain (mesencephalon) of the larval sea lamprey.

Two PmCRH-expressing main hindbrain populations were distinguishable in larvae. The most conspicuous is in the isthmus near the giant isthmic cell and rostrally to the trigeminal motor nucleus (Figures 2g-h, 6g-h). Other neurons were observed in the medial rhombencephalic reticular nucleus, at levels of the facial motor nucleus (Figures 2i, 6i). In P15-P30 prolarvae, two prominent hindbrain populations are clearly separated, one large-celled located in the dorsal isthmus and the other with smaller cells near the ventral midline caudally to the ear vesicle (Figure 9b-d, e), which indicates that they originate in different neural primordia. The caudal cells appear to be the primordium of the PmCRH-expressing reticular cells observed scattered in middle levels of the larval hindbrain. A third hindbrain population of PmCRH-expressing neurons is also observed in prolarvae near the ventral midline of the isthmus. With PmCRH immunohistochemistry, the differences in size and staining intensity between dorsal and ventral isthmic cells were also appreciable (Figure 9e). Similar to adults, in the rostral spinal cord of larvae, a few occasional PmCRH-expressing cells were observed (Figure 7i). No PmCRH-expressing cells were observed in more caudal regions of the spinal cord (at the level of the 5^th^ gill, midbody region or most caudal spinal cord; Supplementary Figure 1c-d).

### 3.2. Distribution of PmCRH-ir fibers in adults and larvae

The use of anti-PmCRH antibody immunofluorescence allowed studying the regional distribution of PmCRH positive fibers in the brain of adults and larvae, together with some observations on the organization of fibers in prolarvae. The density of PmCRH-ir fibers in brains of larvae (around 100 mm in length) and adults varies largely among regions. In the telencephalon of adult lampreys, fine PmCRH-ir fibers are seen in the olfactory bulb, and thicker PmCRH-ir fibers are scattered in the pallium (not shown). In the medial pallium (which developed mainly in adults), a band of fine PmCRH-ir fibers is seen in the neuropil near the periventricular layer of neurons that is characteristic of the lamprey medial pallium. In larvae, the olfactory bulb, pallium and subpallium are mostly devoid of PmCRH-ir fibers.

In the preoptic region and rostral hypothalamus of larvae, the numerous PmCRH-ir fibers coursing in the preoptic-paraventricular-hypophysial tract form the most outstanding tract in the brain (Figure 7a-e), but PmCRH-ir fibers are not present in the caudal hypothalamus and mammillary region. Lateral to the preoptic nucleus, many PmCRH-ir fibers were also appreciable. In adults, these tracts are remarkable, as well as the numerous fibers in the tuberal neuropil (Figure 5a-e). In the larval diencephalon, most fibers course in the neuropil at middle heights (Figure 7f) but lack in the habenula and pineal complex. In the diencephalon of adults, most PmCRH-ir fibers are seen in the thalamus and the prethalamus lateral to the PmCRH-expressing cells, avoiding the optic tract. In addition, the adult pretectum shows abundant PmCRH-ir fibers in the region containing the PmCRH-expressing cells (Figure 5f) and in a positive tract in the transition with the habenula. Some PmCRH-ir fibers cross the midline in the posterior commissure.

In the larval mesencephalon, most fibers run longitudinally in the tegmentum whereas the optic tectum, which is immature in larvae, show scant fibers (Figure 7g). In adults, the optic tectum shows more numerous PmCRH-ir fibers, coursing in the inner cell and fibers layer, but not in outer tectal layers (Figure 5h). In the caudal tectum, however, numerous PmCRH-ir fibers innervate this region, which has some differential features with rostral regions and probably correspond with a specialized superficial region of the torus semicircularis (Sobrido-Cameán et al., 2021b). Abundant PmCRH-ir fibers also cross the midline at this caudal level forming a very conspicuous dorsal commissure (Figure 5i). This PmCRH-ir commissure located between the isthmus and midbrain

In the isthmus of larvae and adults, there are many fibers at most dorso-ventral levels, lacking in the interpeduncular nucleus neuropil (Figure 5k) that extends as a paired ventral neuropil region till the end of the trigeminal motor nucleus (see Yáñez and Anadón, 1994). In the rest of the rhombencephalon, PmCRH-ir fibers are scarce in the dorsal regions (octavolateralis area and dorsal column nucleus) but are abundant in the remainder neuropil regions (Figure 7h). Most PmCRH-ir fibers appear to course longitudinally attending its profile in transverse sections, but at the level of the Mauthner cell (transition between the rhombomeres 3 and 4) there is a small but well-defined commissure of PmCRH-ir fibers crossing as arcuate fibers in the dorsal zone of the raphe. In adults, the pattern is similar (Figure 5k-l), with visceromotor nuclei lacking PmCRH-ir fibers, and scarce fibers in sensory areas (octavolateralis region, dorsal column nucleus and descending trigeminal nucleus).

In P15-P30 prolarvae abundant PmCRH-ir fibers were observed in the marginal layer of basal regions of the brain and spinal cord (Figure 9d-f), most fibers appearing to course longitudinally. In the hindbrain, the isthmic neuropil region was densely innervated by PmCRH-ir fibers since prolarval stages.

### 3.3. Origin of descending PmCRH-ir spinal fibers

In the spinal cord of larvae and adults only a few *PmCRH*-expressing neurons were observed in the lateral horn of the rostral region (close to the obex). But immunofluorescence reveals numerous fibers in the spinal cord at more caudal levels (5^th^ gill, midbody and caudal spinal cord; Supplementary Figure 1a-d) excepting the dorsal column (Supplementary Figure 1a-d) and descending trigeminal root (Figure 7i). In prolarvae, the spinal cord marginal layer also shows PmCRH-ir fibers (Figure 9f). To investigate the brain the origin of these PmCRH-ir fibers, combined PmCRH immunofluorescence and tracing experiments were done in larvae of about 100 mm in length. Combination of retrograde transport of neurobiotin from the spinal cord injected at the level of the 5th gill opening and PmCRH immunohistochemistry revealed retrogradely labeled small reticular hindbrain neurons that were also PmCRH positive (Figure 8), i.e., indicating that these positive cells project to the spinal cord. These cells were located periventricularly mainly in the superior (isthmic) and medial rhombencephalic reticular nuclei near the midline, with conspicuous double labeled cells seen in the isthmus (in the dorsal PmCRH population). A few descending PmCRH cells were occasionally observed in the nucleus of the medial longitudinal fascicle (caudal diencephalon).

### 3.4. Distribution of CRHBP-expressing cell populations in the lamprey central nervous system

We studied the distribution of *PmCRHBP*-expressing cell populations in the brain of the sea lamprey employing specific *PmCRHBP* probes for ISH. The distribution of *PmCRHBP*-expressing cells in lampreys was clearly different from that of PmCRH-expressing perikarya, being mainly located in the septum, striatum, preoptic nucleus, hypothalamus, pineal complex, isthmic/rhombencephalic reticular regions and in the spinal cord. Schematic drawings showing the distribution of *PmCRHBP*-expressing neurons in the brain of adult and larval sea lampreys are presented in Figures 10 and 13, respectively. Photomicrographs of brain sections showing ISH results in adults are presented in Figures 11 and 12, in larvae in Figure 14a-f, and in prolarvae in Figure 14g-i.

**Figure 10.**
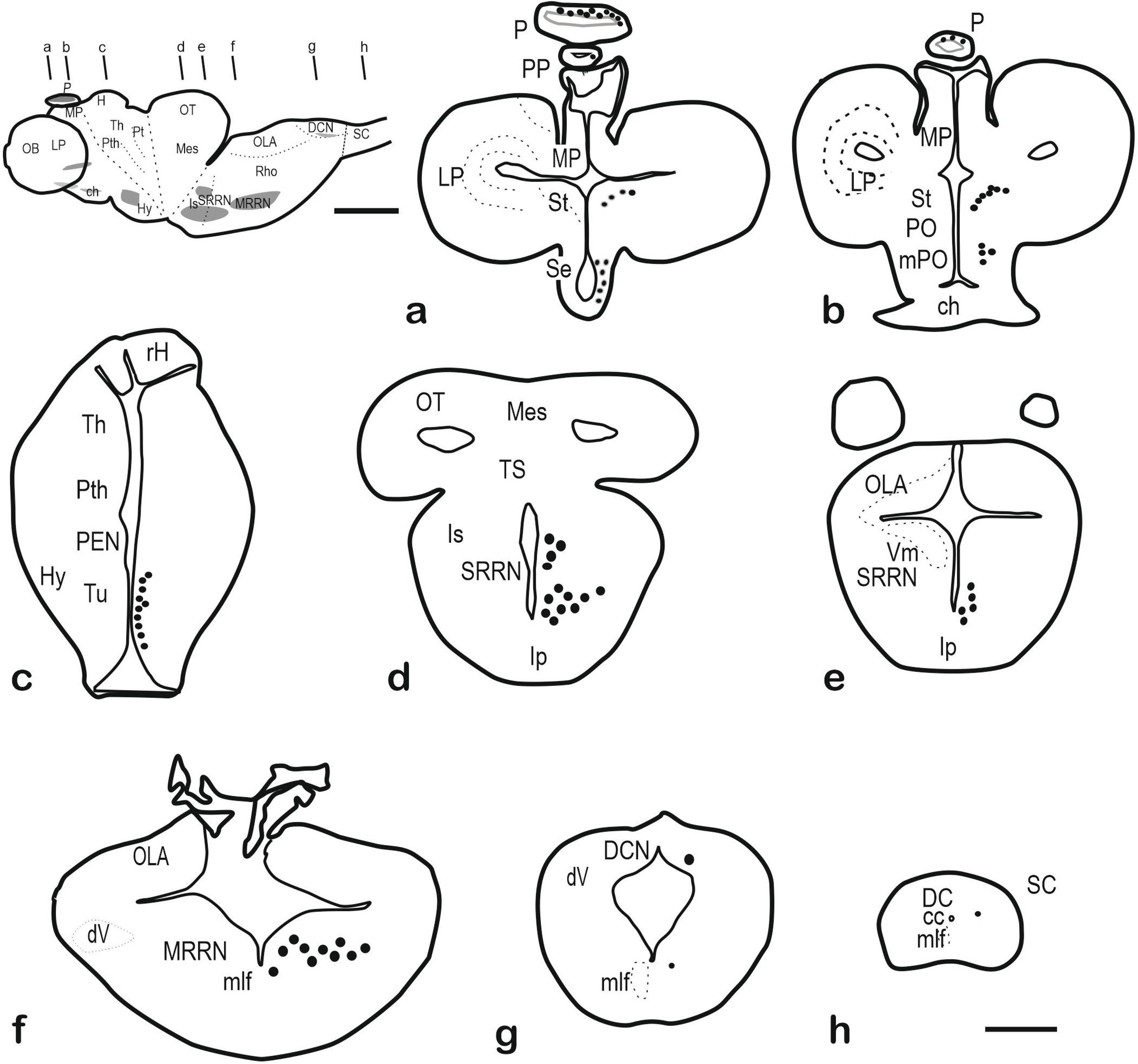
**a-h,** Schematic drawings of transverse sections of the brain of an adult sea lamprey showing the distribution of *PmCRHBP-*expressing neurons (at the right) and the anatomical structures (at left). The levels of sections are indicated in the figurine of the lateral view of the brain. For abbreviations, see the list. Scale bars, 200 µm (a-h), 1 mm (figurine).

**Figure 11.**
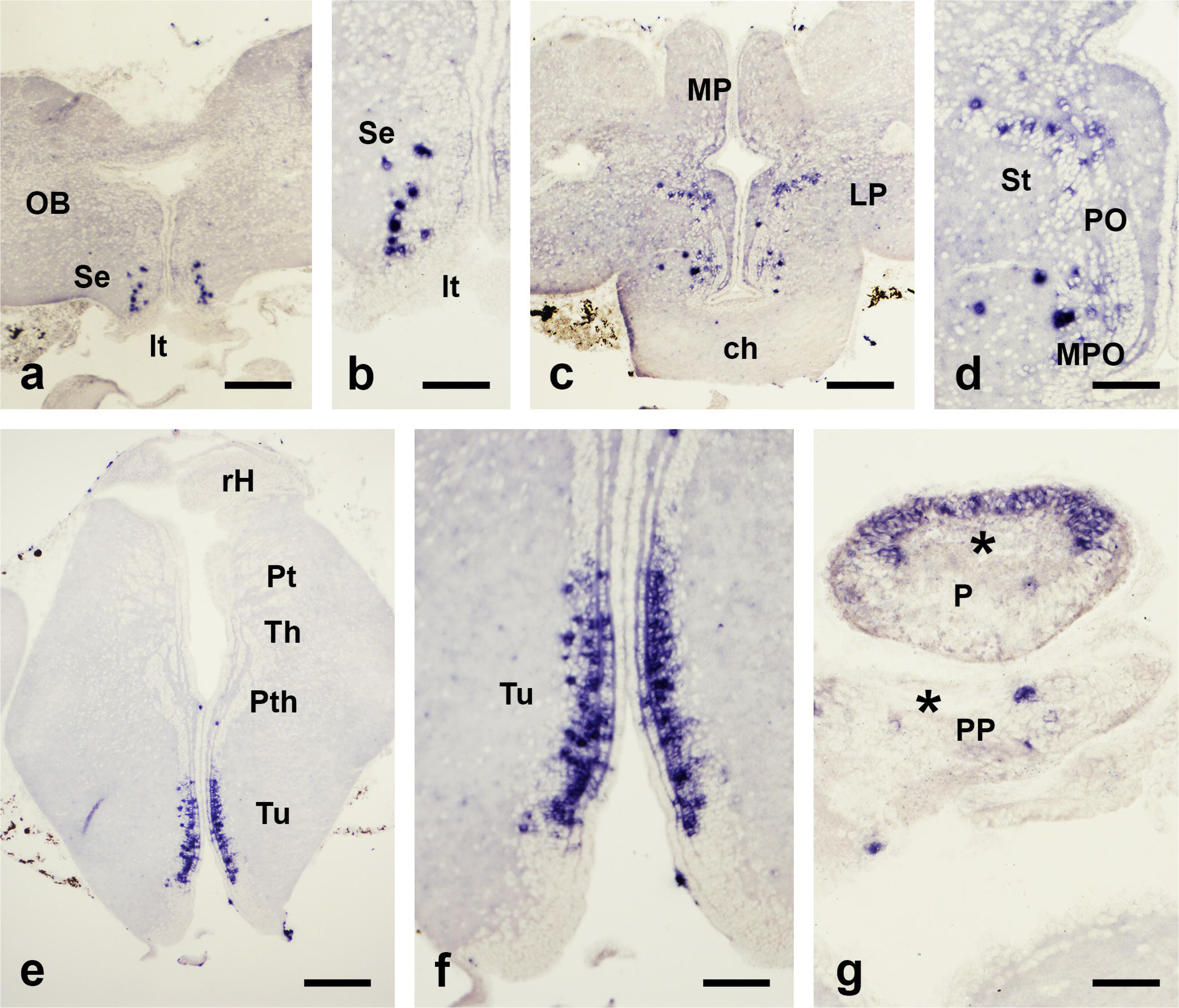
Photomicrographs of transverse sections of the forebrain of a young sea lamprey showing neurons expressing *PmCRHBP* mRNA revealed by ISH. **a-b**, Panoramic view and detail of the olfactory bulb and septal-terminal lamina region showing numerous positive neurons in the septum. **c-d**, Panoramic view and detail of the telencephalon and preoptic region showing PmCRHBP-expressing neurons in the striatum and outer zone of the mPO nucleus. **e-f**, Panoramic view of the hypothalamus and diencephalon and detail of the tuberal nucleus population of positive neurons. **g**, Section through the epithalamus showing positive cells in the outer wall of the pineal organ (P) and a few cells in the parapineal organ (PP). Asterisks, vesicle lumen. For abbreviations, see the list. Scale bars, 50 µm (b, d, f, g), 125 µm (a, c, e).

**Figure 12.**
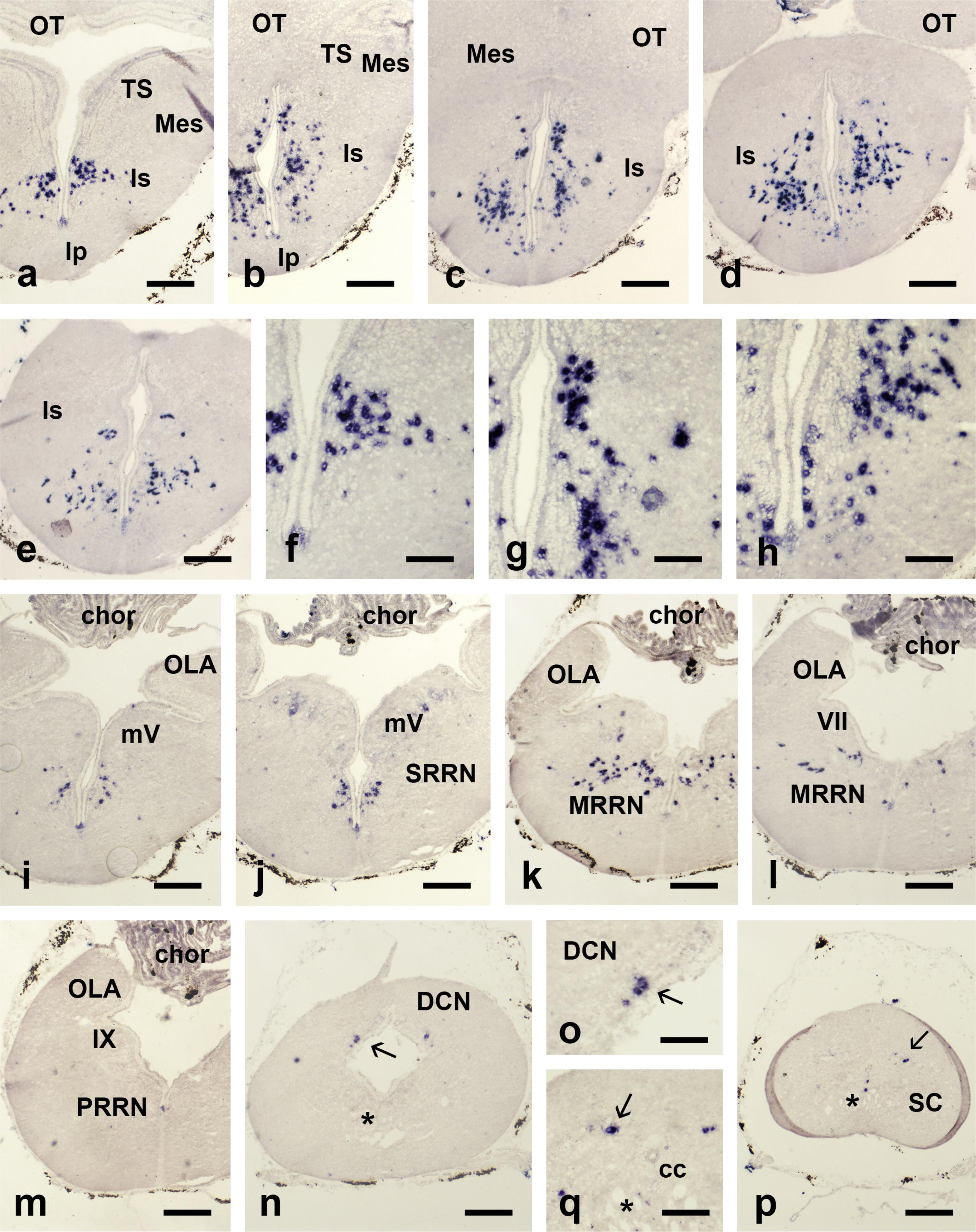
Photomicrographs of transverse sections of the hindbrain of a young sea lamprey showing neurons expressing *PmCRHBP* mRNA revealed by ISH. **a-e**, Panoramic views of the isthmus region showing abundance of *PmCRHBP* positive neurons. Note that the isthmus region appears below the mesencephalon (which is free of CRHBP-expressing cells) at rostral levels. **f-h**, Details of positive cells in rostral, intermediate and caudal isthmic levels. **i-m**, Panoramic views of sections of the hindbrain between rhombomere 2 and 6 showing differences in distribution *PmCRHBP*-expressing neurons along the medulla: i-j, level of the trigeminal motor nucleus (rhombomeres 2-3); k-l, levels of the octaval nerve entrance and facial motor nucleus (rhombomeres 4-5); m, level of the motor glossopharyngeal nucleus (rhombomere 6). **n-o**, Section and detail at the level of transition of the hindbrain to the spinal cord, caudal to the calamus, showing positive neurons (arrowed) in the dorsal column nucleus. **p-q**, Panoramic view and detail of the spinal cord showing a few PmCRHBP positive neurons (arrows). Asterisks in **n**, **p** and **q** indicate the medial longitudinal fasciculus. For abbreviations, see the list. Scale bars, 125 µm (a, b, c, d, e, i, j, k, l, m, n, p), 50 µm (f, g, h, o, q).

**Figure 13.**
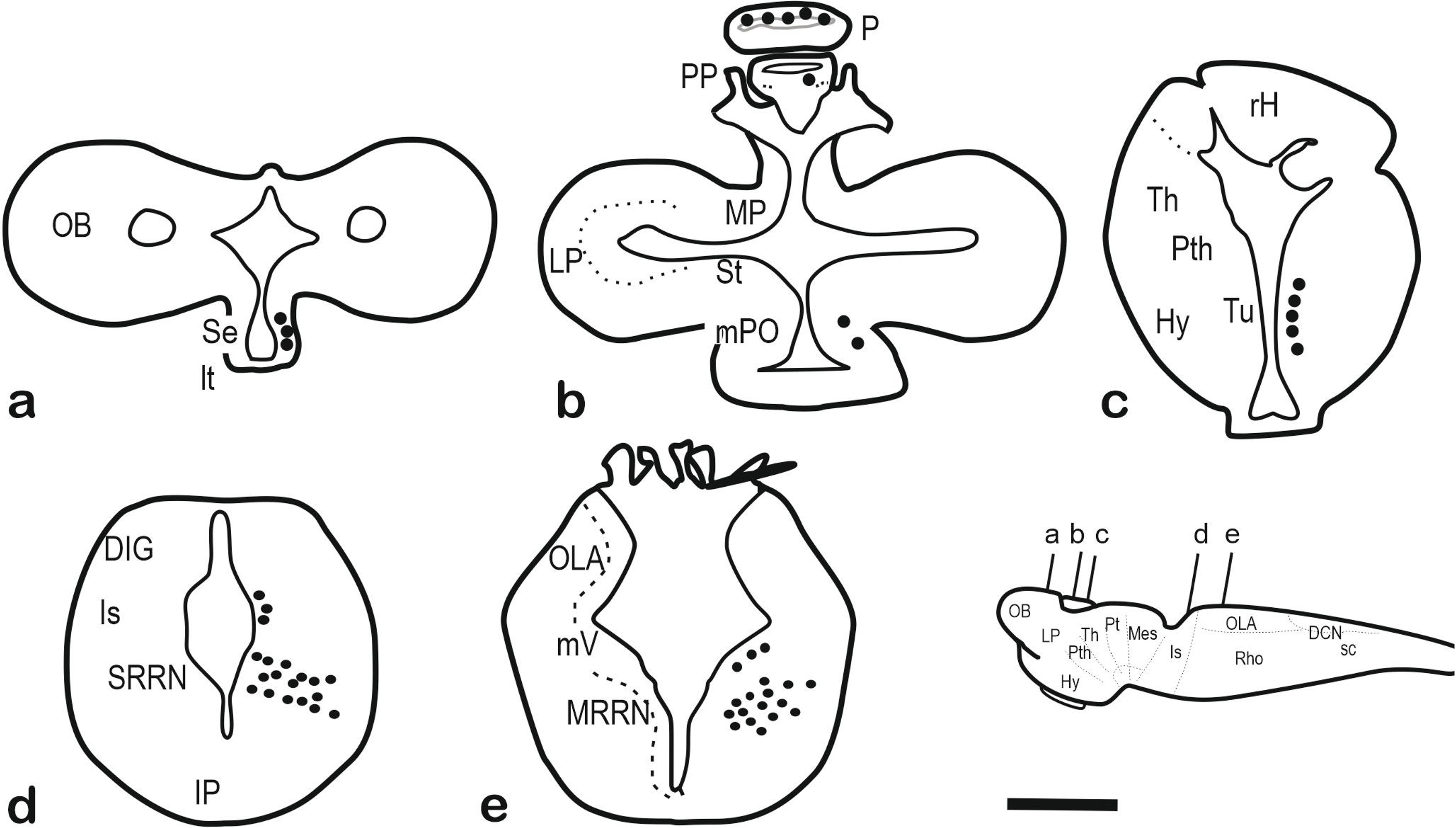
**a-e**, Schematic drawings of transverse sections of the brain of a sea lamprey larva (about 100 mm long) showing the distribution of *PmCRHBP-*expressing neurons (at the right) and the anatomical references (at left). The levels of sections are indicated in the figurine of the lateral view of the brain. For abbreviations, see the list. Scale bar, 200 µm (a-e).

**Figure 14.**
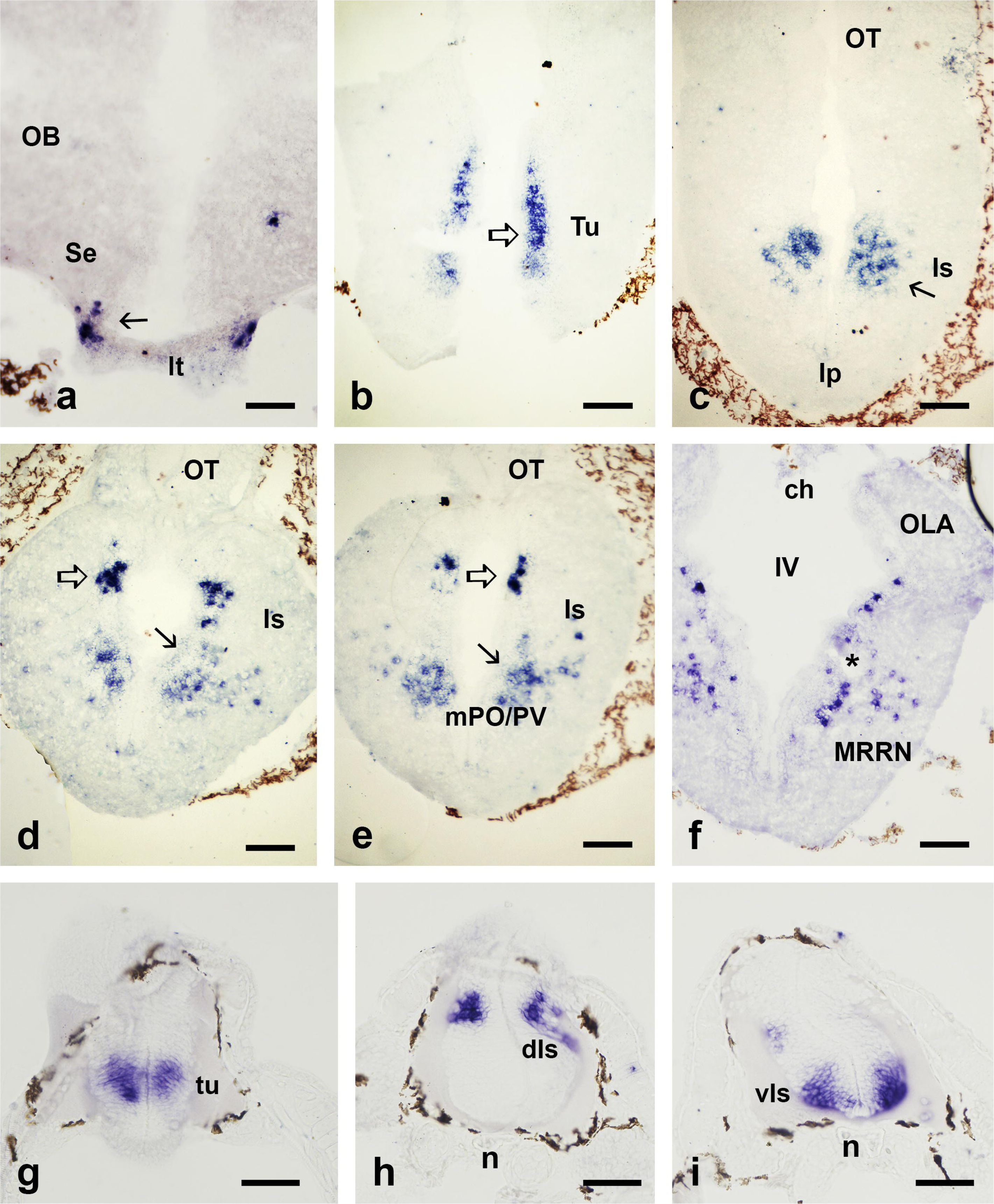
Photomicrographs of transverse sections of the brain of a larval sea lamprey (100 mm long) (a-f) and of a P15 prolarvae (g-i) showing neurons expressing *PmCRHBP* mRNA revealed by ISH. **a**, Section through the telencephalon at the level of the lamina terminal showing positive neurons in the septum (arrow). **b**, Section through the hypothalamus showing a periventricular band of positive cells in the tuberal nucleus (outlined arrow). **c-e**, Sections through the isthmus region (from rostral to caudal) showing groups of dorsal (outlined arrows in d-e) and ventral (arrows) positive neurons with different appearances. **f**, Section at the level of the octaval nerve entrance showing abundant positive neurons in the medial rhombencephalic reticular nucleus. Asterisk in f indicates a large reticulospinal neuron. **g**, Section through the prosencephalon of a P15 prolarvae showing a dense population of *PmCRHBP*-expressing neurons corresponding to the tuberal nucleus. **h-i**, sections through the isthmus showing groups of *PmCRHBP*-expressing neurons. For abbreviations, see the list. Scale bars, 50 µm.

In young adults, the most rostral *PmCRHBP*-expressing cells were observed in the walls of the septum (Figures 10a, 11a-b). This population extended to the preoptic nucleus with cells scattered in a location lateral to the main band of PmCRH-expressing cells (mPO/paraventricular nucleus) (Figures 10b, 11c-d). Both cell populations (*PmCRHBP*- and PmCRH-expressing) scarcely intermingle in the preoptic region, as shown by double PmCRH immunohistochemistry / *PmCRHBP* ISH (not shown). In addition, a small population of *PmCRHBP*-expressing cells was observed in the striatum, located in the characteristic cell band and extending toward the periventricular zone of the ventral pallium (Figures 10a-b, 11c-d). In the hypothalamus (anterior tuber and tuberal region), a long band of *PmCRHBP*-expressing cells was observed in intermediate-dorsal region from the nucleus of the postoptic commissure till the rostral level of the hypophysis (Figures 10c, 11e-f). Most cells are located within the dense rows of cells parallel and away from the ependyma, but a few cells are located among ependymal cells. Again, this location is different from that of PmCRH-expressing cells of the tuberal region, which located in the most ventral region (see above). In the diencephalic roof, numerous *PmCRHBP*-expressing cells were observed in the walls of the parapineal and pineal organs (Figures 10a-b, 11g). In the pineal vesicle, most of positive cells are distributed in the dorsal (outer) wall, whereas in the parapineal vesicle they mostly are in a lateroventral region.

In the isthmus-first rhombomere of adults (which lies rostral to the trigeminal motor nucleus), different populations of *PmCRHBP*-expressing cells were observed (Figures 10d-e, 12a-h). The most dorsal was a rather compact group of cells located in the periventricular layer of cells at an intermediate dorso-ventral level close to the giant I1 cell. A second group, much more numerous than the former, occupies a more ventral location and is clearly differentiated by the scattered distribution of many cells away the periventricular layer through a lateral wing-like area. In this ventral group, smaller sized and paler cells were also observed close to the ventral ependymal layer. More scattered populations of *PmCRHBP* positive cells were observed in the rhombencephalon caudal to the isthmus-first rhombomere till the rostral spinal cord (Figures 10f-g, 12j). These cells are located in basal plate derived regions medial to the visceromotor nuclei and in the reticular formation close to the medial longitudinal fascicle. These cells are scarce at levels of the trigeminal motor nucleus (rhombomeres 2-3), increase clearly in number at levels of the facial motor nucleus (rhombomeres 5-6) (Figures 10f, 12i-l), and disappear in more caudal hindbrain levels (Figure 12m). At the transition with spinal levels, some *PmCRHBP* positive cells were also observed in the nucleus of the dorsal column (Figures 10g, 12n-o). In the rostral spinal cord, *PmCRHBP*-expressing cells are observed from the hindbrain-spinal transition, with a few cells per section located below the central canal and in the lateral horn (gray matter) (Figures 10h, 12p-q).

In larvae of about 100 mm in length, some *PmCRHBP*-expressing cells were observed in the septum (Figures 13a, 14a), and occasional cells in the preoptic nucleus (Figure 13b). The hypothalamic population of *PmCRHBP*-expressing cells was numerous in the tuberal region (Figures 13c, 14b). In the pineal complex, *PmCRHBP*-expressing cells were observed in the walls of the parapineal and pineal vesicles (Figure 13b). Two isthmic groups were recognizable in these larvae (Figures 13d, 14c-e). The dorsal isthmic *PmCRHBP*-expressing cells are located near the I1 giant neuron, and the ventral population is more scattered and extended laterally. In addition, scattered *PmCRHBP*-expressing cells were observed in the rhombencephalic reticular region caudal to the trigeminal nerve entrance till caudal levels (Figures 13e, 14f). In the rostral spinal cord, scarce positive cells are scattered in the gray matter without a defined location.

No positive cells were observed in the preoptic region or in the septal primordium of prolarvae. In P15-P25 prolarvae, three well defined groups of *PmCRHBP*-expressing cells were observed in the mantle layer of the brain, and a few positive small cells were appreciable in the pineal vesicle. The most rostral *PmCRHBP* positive group consists of a compact cell cluster in the cell mantle that is located in the primordium of the hypothalamus at intermediate dorso-ventral levels (Figure 14g). This level is recognizable in transverse sections because it is caudal to the postoptic commissure, being also distinctive that ventrally the brain is close to the preoral epithelium without intervening notochord. At the level of the isthmus/first rhombomere (see Meléndez-Ferro et al., 2003), two groups of *PmCRHBP*-expressing neurons were already present in prolarvae, one more rostral and in intermediate dorso-ventral level, and the other located in the ventral region of the mantle layer slightly caudal to the former (Figure 14h-i). In head sections at this level the notochord tissue was clearly appreciable between the brain and the preoral epithelium. Caudal to these isthmic groups, *PmCRHBP*-expressing cells were observed in the most ventral hindbrain/spinal region, laterally to the small floor plate. In these prolarvae, the limits between the hindbrain and the rostral spinal cord are difficult to assess, but some of these cells appear to be in spinal levels.

## 4. Discussion

In this study we analyzed in several life stages of the sea lamprey the distribution of *PmCRH*- and *PmCRHBP*-expressing cells using ISH with probes specific for these mRNAs. For PmCRH, too, we generated a specific antibody (as shown by ELISA experiments) raised against a synthetic peptide with the putative sequence of the mature PmCRH peptide. Complementary experiments included double stainings (PmCRH immunohistochemistry with TH immunohistochemistry or *PmCRHBP* ISH), as well as experiments with a combination of PmCRH immunohistochemistry with retrograde transport of neurobiotin from the spinal cord. Previous studies showed expression of CRH and CRHBP mRNA in extracts of the whole sea lamprey brain at different life stages by RT-qPCR (Endsin et al., 2017), but not its neuronal distribution as shown here. As far as we are aware, our results present the first report about the organization of CRH- and CRHBP-expressing neurons in the brain of any jawless vertebrate.

Regarding the distribution of neurons in the sea lamprey brain and rostral spinal cord, our results reveal that *PmCRH*- and *PmCRHBP*-expressing cells represent separate populations with distinct distributions, and thus this neuropeptide and its binding protein are probably synthesized and released by two different systems. Note, however, that the single CRHBP of mammals binds both CRH and Ucn1 with sub-nanomolar affinity (Ketchesin et al., 2017a), and in Xenopus CRHBP binds with high affinity to CRH-like peptides (Valverde et al., 2001). Thus, it is probable that PmCRHBP binds to several members of the CRH family of lampreys. The brain distribution of the mRNA of another member of the CRH gene family (Ucn3) has been recently studied by our group (Sobrido-Cameán et al., 2021a). Comparison of Ucn3 distribution results in sea lamprey with present PmCRH distribution reveals important differences between the distributions of these substances. Since we could only study the distribution of PmCRH fibers, the relation between the CRHergic cells with that of Ucn3- or PmCRHBP-expressing axons was not determined. Moreover, the mode of secretion of CRHBP by neurons in vertebrates (constitutive or regulated) is not yet clear.

The lamprey CRHergic preopto-paraventricular-neurohypophysial system was outstanding in larvae and adult lampreys, but not in P15 prolarvae. Many neurons belonging to the lamprey mPO and its caudal continuation as the paraventricular nucleus express *PmCRH* mRNA in both larvae and adults. PmCRH immunohistochemistry shows that cells of this continuous cell population send axons toward the tuberal hypothalamus and reach the neurohypophysis since prolarval stages, where presumably form terminal endings on the meningeal surface. Recently, CRH-ir neurons have been reported with immunohistochemistry in the preoptic nucleus and periventricular preoptic nucleus of a hagfish (Amano et al., 2016). This preoptic CRH neurosecretory system appears well conserved in teleost fishes (rainbow trout: Ando et al., 1999; zebrafish: Alderman and Bernier, 2007; Japanese eel: Amano et al., 2014), although in mammals (and probably also in reptiles and birds) the homologous nuclei are two separate entities, the supraoptic and the paraventricular nuclei, both with neurons producing CRH and sending axons to the hypophysis (Keegan et al., 1994; Wamsteeker Cusulin et al., 2013). Unlike in mammals, the preopto-paraventricular CRHergic neurons of sea lamprey (at least in part) appear to be of CSF-contacting type, a type of neuron common in the lamprey hypothalamus (Barreiro-Iglesias et al., 2017).

The release of CRH by fibers of the hypothalamo-hypophysial tract in the lamprey neurohypophysis is thought to promote the secretion of adrenocorticotropin (ACTH) by the adenohypophysis, although there are no direct experimental results supporting this hypothesis (Roberts et al., 2014). In the sea lamprey and other lampreys, two different hormone precursors of the pituitary hormones melanotropin (MCH) and ACTH have been identified, proopiomelanotropin and proopiocortin, respectively (Ficele et al., 1998). This is unlike the single common precursor proopiomelanocortin gene present in most gnathostomes, excepting in various teleosts in which this gene is duplicated (Alsop and Vijayan, 2009). In the sea lamprey and other lamprey species, these precursors, as their derived peptides MSH and ACTH, are expressed in cells of different parts of the adenohypophysis (pars intermedia and pars proximalis), both in larvae and adults, although with high quantitative variation between larvae, transforming lampreys and different adult phases (Nozaki et al., 1995, 2008; Ficele et al., 1998; Takahashi et al., 2006). Older studies revealed that both stress and ACTH injection affect to cells of the interrenal tissue (adrenocortical homologue) of sea lampreys (Hardisty, 1972; Youson, 1972). Thus, the hypothalamo-hypophysial-interrenal (adrenal) axis appears to be conserved between agnathans and gnathostomes, indicating that it appeared in common ancestors of all vertebrates. Paradoxically, the lamprey interrenal tissue does not synthesize the stress hormones cortisol or corticosterone and thus the function of this tissue as a producer of stress hormones is unclear (Roberts et al., 2014). A steroid molecule found in blood (11-deoxycortisol) has been proposed as the corticosteroid hormone of lampreys, although its regulation and sites of synthesis are not yet known (Close et al., 2010). Recent data indicate that 11-deoxycortisol is involved in hydromineral balance (Shaughnessy et al., 2020) and is elevated by stress, activating gluconeogenesis (Shaughnessy and McCormick, 2021). A study in a hagfish, however, has detected cortisol and corticosterone in plasma (Amano et al., 2016), which is an intriguing difference with their reported absence in lampreys.

Neurogenetic studies in mice have determined that the CRH-expressing neurosecretory neurons of the paraventricular nucleus appear in a primordial embryonic brain region (supraopto-paraventricular primordium) expressing the gene Orthopedia (*otp*) (Acampora et al., 1999; Morales-Delgado et al., 2014). This transcription factor is also essential for restricting the fate of other categories of neuroendocrine neurons (vasotocinergic, TRHergic: Acampora et al., 1999). Knocking-out the *otp* gene in mice embryos precludes the appearance of CRH-expressing populations, indicating a dependence on this transcription factor in the specification of cells of this region (Acampora et al., 1999; Morales-Delgado et al., 2014). In adult mice, the CRH-expressing paraventricular population is activated by stress (Wamsteeker Cusulin et al., 2013). *otp* is also expressed in the corresponding band that gives rise to the preoptic CRH population in zebrafish (Herget et al., 2014). This subdivision, named in mammals as the paraventricular or supraoptoparaventricular area in prosomeric models is currently ascribed to the alar region of the hypothalamus (Morales-Delgado et al., 2014). An alternative model suggests it is part of an optic related region that is outstanding in fishes (Yamamoto et al., 2017). Although the expression of *otp* is not known in the lamprey brain, most probably the PmCRH-expressing population of the magnocellular preopto-paraventricular band is homologous to that of teleosts and mammals. This region also contains abundant vasotocinergic and TRHergic neurons in lampreys (Goossens et al., 1977; Pombal and Puelles, 1999; De Andrés et al., 2002). Our results with double staining with *PmCRH* ISH and TH immunohistochemistry further reveal the mutual exclusion of cellular territories of PmCRH and TH expression in dorsal and ventral zones of the periventricular region of the lateral optic recess, respectively.

An additional PmCRH-expressing hypothalamic population of small cells appears in the ventral tuberal region of the hypothalamus in adults, although not in larval lampreys. The significance of this tuberal population and the possibility that these cells may be related with the hypophysis were not determined. The presence of CRHergic hypothalamic populations outside the preopto-paraventricular region was also reported in the so-called ventral hypothalamus of *Lepisosteus* and zebrafish and in the nucleus lateralis tuberis of *Oreochromis mossambicus* (tilapia) (Alderman and Bernier, 2007; Grone and Maruska, 2015), which might correspond to those of the lamprey ventral tuberal region. Whereas in *Lepisosteus* and in tilapia only a CRH gene (CRH1 or CRHb, respectively) is expressed there, in zebrafish both CRHa and CRHb are expressed.

Several *PmCRH*-expressing populations unrelated with the hypophysis were observed in the lamprey brain, some of them appearing early in prolarvae (in the isthmus or caudal hindbrain), and others appearing at more advanced stages (large larvae or adults). Among the late appearing *PmCRH*-expressing populations are those found in the telencephalon (olfactory bulb and pallio-striatal populations). In the olfactory bulb, *PmCRH*-expressing neurons are very scant in larvae and more numerous in adults. In adults, immunohistochemistry also shows very thin PmCRH-ir fibers branching in the olfactory bulb granular layer but not, or scarcely, in olfactory glomeruli. This suggests that these neurons are not typical granule cells involved in local circuits of the olfactory bulb. No cells or CRH-ir fibers were observed in the olfactory bulb of a hagfish (Amano et al., 2016), which is unlike that observed here in the sea lamprey. However, CRH-ir neurons were reported in the glomerular layer of the olfactory bulb of the teleost fish *Oreochromis mossambicus* (Pepels et al., 2002) and of mammals, where they appear in the external plexiform layer (Garcia et al., 2016; Peng et al., 2017; Wang et al., 2021). Studies in mice have shown that these neurons act in local circuits via release of CRH, being essential for shaping olfactory circuits (García et al., 2016). In mouse olfactory bulb, CRH expression first appears postnatally (García et al., 2016), whereas our results suggest late expression in lamprey larvae. Anyway, correspondences between CRHergic cell types of the olfactory bulb of lampreys and mammalians need further clarification.

With regard the telencephalic lobes, a sparse population of PmCRH-expressing cells extends between the striatum and ventral pallium, but it is only present in adult lampreys (both young and mature). Again, this suggests that they are related to some functions specific to adult life. Interestingly, this CRHergic population is topographically closely related with a population of cells expressing *PmCRHBP* in adults, suggesting functional relationship between these two populations. This codistribution reminds the populations of CRH- and CRHBP-expressing cells reported in the ventral nucleus of the zebrafish subpallium (Alderman and Bernier, 2007). In tilapia, however, the largest CRHergic population in the brain was observed in the lateral nucleus of the subpallium (Pepels et al., 2002). By its location, the lamprey striato-pallial PmCRH-expressing population may remind the CRHergic neurons reported in the amygdala of mammals (Potter et al., 1992; Peng et al., 2017), which is involved in stress control, or to caudo-putamen neurons (Wang et al., 2021), but functions of the lamprey striatum in stress control are unknown. Also, the absence of proper pallial CRHergic cells in the sea lamprey reveals important differences with mammals, which show abundant CRHergic neurons in some cortical regions (Potter et al., 1992; Garcia et al., 2016; Peng et al., 2017; Wang et al., 2021). Furthermore, ICH reveals that most of the pallium of lampreys (dorsal, lateral) shows scant CRHergic innervation, although the adult medial pallium of adults shows a more abundant innervation near the periventricular region. In the teleost fish tilapia, very numerous CRH-ir fibers were observed in the rostral pole of the pallium, possibly coming from a subpallial (Vl) population of CRH-ir neurons (Pepels et al., 2002). Although a subpallial (striatal) population of PmCRH-expressing neurons is present in adult sea lamprey, no territory densely innervated by PmCRH-ir fibers as that reported in tilapia was observed in the pallium.

The lamprey prethalamus (alar diencephalon) shows populations of PmCRH-expressing neurons in adults, but not in larval stages. A small pretectal PmCRH-expressing group observed in adults and large larvae is located in the same pretectal region of a small pretecto-habenular group reported with tracing methods (Yáñez et al., 1999; Stephenson-Jones et al., 2012), suggesting that PmCRH is involved in these habenular circuits. However, PmCRH immunohistochemistry only shows scant PmCRH-ir fibers in the habenula and parapineal ganglion. The most abundant PmCRH-expressing diencephalic population is the prethalamic one, which is distributed sparsely close to the dorsal (caudal) hypothalamus, whereas the other is found in the pretectum. Cells of the lamprey prethalamus (formerly ventral thalamus) project to the hindbrain and spinal cord, and this region is considered as premotor (El Manira et al., 1997). However, we did not find PmCRH-expressing prethalamic cells with spinal projections in our combined tracing experiments. A third diencephalic PmCRH-expressing population is found in the nucleus of the medial longitudinal fascicle. Cells of this nucleus project to the rhombencephalic reticular formation (Capantini et al., 2017), and fibers from PmCRH-expressing cells of this region, together with those originated in the isthmic and hindbrain reticular cells, might form part of longitudinal tracts observed with immunohistochemistry.

Regarding the absence of PmCRH-expressing cells in the lamprey mesencephalon, this is in contrast with the presence of CRH-expressing cells in the midbrain of tilapia (periventricular layer of the optic tectum; Pepels et al., 2002). Anyway, the observation of dense innervation by CRHergic fibers in the caudal midbrain tectum of adult sea lampreys is interesting, as well as the conspicuous caudal commissure. This caudal tectal region may represent in fact a specialized portion of the torus semicircularis, as suggested on the base of studies of distribution of glycine (Villar-Cerviño et al., 2008) and Ucn3 (Sobrido-Cameán et al., 2021a). In addition, this caudal region is innervated by afferents from the octavolateralis region (De Arriba Pérez, 2007), as the topologically similar torus semicircularis of *Xenopus* located just caudal to the optic tectum in the tectal lamina (Morona and González, 2009). Other midbrain regions also show rich innervation by CRHergic fibers, but whether they are en-passant fibers that contact midbrain neurons was not established. With regard the binding protein, no *PmCRHBP*-expressing cell was observed in the lamprey midbrain, and innervation by positive fibers from other areas is not known.

The lamprey isthmus/first rhombomere region contains abundant PmCRH-expressing neurons since the earliest developmental stages investigated, which originate in two different clusters, dorsal and intermediate-ventral. Even in prolarvae, the morphology of cells is different in the dorsal and intermediate-ventral populations, and these differences are also appreciable in adults. Interestingly, the isthmus/first rhombomere also contains the largest population of *PmCRHBP*-expressing cells, which also appears as two well separated cell clusters in prolarvae. Our results suggest a close developmental, topographical and functional relation between these PmCRH- and *PmCRHBP*-expressing isthmic/superior rhombencephalic populations. This lamprey isthmic region is an important functional hindbrain center that controls locomotion, although more often it is called as “mesencephalic locomotor region” (MLR) (Sirota et al., 2000; Brocard and Dubuc, 2003; Grätsch et al., 2019), which leads to anatomical confusion (see Villar-Cerviño et al., 2008). Comparison of maps of cells in “MLR” levels presented in Grätsch et al. (2019) with those of present results also confirms that most of these cells are clearly isthmic, not mesencephalic. Physiological studies reveal that projections from neurons of this region to neurons of the middle rhombencephalic reticular nucleus can activate or stop locomotion via glutamatergic transmission (Grätsch et al., 2019). The abundance of PmCRH- and *PmCRHBP*-expressing cells of this region suggests that these neurons could be involved in stress control of locomotion. The isthmus/first rhombomere of the sea lamprey shows abundant serotonergic cells from prolarvae to adulthood (Abalo et al., 2007; Barreiro-Iglesias et al., 2008a; Cornide-Petronio et al., 2013), with groups ventral, intermediate and dorsal, which in part are codistributed with PmCRH- and *PmCRHBP*-expressing neurons. The possibility of colocation of serotonin with PmCRH or PmCRHBP in these cells was not investigated. In the mammalian isthmus, an important population of CRH-expressing cells is found in the Barrington’s nucleus, which is close to the locus coeruleus containing noradrenergic neurons. The Barrington’s nucleus is an important micturition center in rodents projecting axons to the lumbosacral spinal cord (Valentino et al., 2000; Verstegen et al., 2017), being part of a sympatho-adrenomedullary stress system (Chaves et al., 2021). However, lampreys have no bladder like that of mammals, and probably their isthmic CRHergic populations are involved in other roles. Hindbrain levels contain other PmCRH-expressing neurons generally in reticular areas. In mammals, CRHergic cells are also observed in the parabrachial nucleus, the tegmental nucleus, ventrolateral medulla, medial octaval nucleus and inferior olivary complex (Peng et al., 2017; Chaves et al., 2021). However, correspondence with lamprey hindbrain PmCRH-expressing populations is unclear. Moreover, the sea lamprey lacks catecholaminergic populations in the rostral rhombencephalon and does not have a locus coeruleus homologue (Barreiro-Iglesias et al., 2010b), and lacks an inferior olivary nucleus, whose CRHergic cells project to the cerebellum in mammals (which is also absent in lampreys; see Lammanna et al., 2022).

Prompted by the abundance of PmCRH-ir fibers coursing longitudinally along the brain and spinal cord, we performed some experiments of neural tracing with neurobiotin combined with PmCRH immunofluorescence. Our results show double labeled CRHergic cells in the superior and medial rhombencephalic reticular nuclei and occasionally in the diencephalon (nucleus of the medial longitudinal fascicle). Descending fibers are probably early appearing, judging from the abundance of CRHergic fibers in the prolarval spinal cord, but target cells are not known.

### 4.1 Early appearance of CRH- and CRHBP-expressing hindbrain populations

The presence of conspicuous PmCRH- and PmCRHBP-expressing cells in the rostral hindbrain was noted in the brains of P15 prolarvae, the first developmental stage studied, suggesting that these populations appeared in earlier stages. The hindbrain PmCRH-expressing populations appear before than those in the preoptic region, which is in line with gradients of maturation observed in the brain of lamprey for other neurotransmitters as GABA and serotonin (Meléndez-Ferro et al., 2002a, 2003; Abalo et al., 2007). The precedence of hindbrain PmCRH-expressing populations in lamprey brain is in contrast with observations in the zebrafish, in which the preoptic neurons are those appearing first (Chandrasekar et al., 2007). Also, the delayed appearance of striatal PmCRH-expressing neurons in lampreys till juvenile stages contrasts with the earlier appearance of subpallial, preoptic and posterior tubercle CRH neurons than hindbrain neurons in zebrafish embryos (Chandrasekar et al., 2007). This comparison reveals marked heterochrony between zebrafish and lampreys, which would be related to the extended larval life of lampreys showing major differences with the adult phases. Differential regulation in CRH neuronal populations was also reported in mouse embryos (Keegan et al., 1994), with cells in the paraventricular nucleus, Barrington’s nucleus, olivary complex, and amygdaloid primordia appearing on embryonic day 13.5, whereas CRH mRNA is not detectable in the cortex until after birth.

### 4.2. Comparison of CRHBP and CRH distributions

CRHBP is a phylogenetically ancient peptide whose presence has been demonstrated in all major vertebrate groups from fish to mammals (Seasholtz et al., 2002; Endsin, 2013), although its brain distribution has been only shown in mammals (Potter et al., 1992). In mammals, this protein binds with high affinity equally to CRH and UCN1 and sequesters these ligands away from the receptor; this modifies the action of these factors on their receptor with a variety of effects on various targets (for review see Seasholtz et al., 2002). We showed for the first time the distribution of *PmCRHBP* in the brain of a lamprey. Our results show a limited brain distribution of *PmCRHBP*-expressing cells in adults, with positive neurons mainly located in the septum, striatum, preoptic nucleus, dorsal hypothalamus, pineal and parapineal organs, dorsal and ventral isthmus, and some additional rhombencephalic populations. Only some of these areas with *PmCRHBP*-expressing neurons also show PmCRH-expressing cells (striatum, preoptic-paraventricular nucleus and isthmus). On the other hand, studies in the rat (Potter et al., 1992) show huge numbers of CRHBP positive cells in forebrain regions that lack these cells in the sea lamprey, such as the olfactory bulb, cortex (general cortex, pyriform cortex, and hippocampus), amygdaloid complex, etc. Also, important CRHBP expression is found in both the superior and inferior collicules, and in the principle trigeminal sensory nucleus (Potter et al., 1992), which in lamprey are free of *PmCRHBP* positive neurons. These authors have shown that most cells in nuclei with expression of both substances do not colocalize them but are expressed by different neurons. This appears to be the case in most lamprey brain regions, but the possibility that both proteins were co-expressed by some cells in regions showing both substances (as in the isthmus) cannot be ruled out. With regards the nature of CRHBP-expressing cells in the mammalian prefrontal cortex, some studies indicate that they are preferentially GABAergic neurons that also express somatostatin (Ketchesin et al., 2017b). Comparison of distribution of *PmCRHBP* with those of cells expressing each of the three somatostatin genes reported in lamprey (Sobrido-Cameán et al., 2021b), however, does not show close correspondence with any of them. From a comparative point of view, the lack of similar studies in other vertebrates precludes to propose a phylogenetic hypothesis that would be based on mere speculation.

### 4.3. *CRHBP* in the pineal complex

Lampreys show a pineal complex that consists of a pineal organ (vesicle) and a parapineal organ (vesicle plus ganglion), which show different organization, maturation patterns and brain connections (see Pombal et al., 1999; Yáñez et al., 1999; Meléndez-Ferro et al., 2002b; Barreiro-Iglesias et al., 2017). *PmCRHBP*, but not PmCRH, is expressed in cells of the pineal vesicle of the sea lamprey since prolarval stages. These cells correspond by location to dorsal photoreceptors that express alpha-transducin, beta-arrestin and parapinopsin but not to those ventral photoreceptors expressing rhodopsin (Kawano-Yamashita et al., 2015; Barreiro-Iglesias et al., 2017). This suggests PmCRHBP expression and release by parapinopsin pineal photoreceptors that are sensible to color of light. Axonal projections from long-axon pineal photoreceptors to the pretectum and optic tectum have been reported in lampreys (Pombal et al., 1999), and thus the pineal organ might send *PmCRHBP*-expressing projections to the dorsal diencephalon and midbrain. Neural cells of the parapineal vesicle only project to the adjacent parapineal ganglion (Yáñez et al., 1999), which is a specialized region of the left habenula, and thus probably these cells exert there a local function. Interestingly, a small pretectal nucleus projects to the parapineal ganglion and habenula (Yáñez et al., 1999; Stephenson-Jones et al., 2012). By location and morphology, this population strongly reminds the PmCRH population in the same region. Whether the PmCRH-expressing pretectal cells are the same that send axons to the parapineal ganglion and habenula should be assessed in the future.

### 4.4. Conclusions

Present results reveal for the first time the location and organization of CRH- and CRHBP-expressing neurons in the brain of a jawless vertebrate. Our results show that, in the sea lamprey, *PmCRH*- and *PmCRHBP*-expressing cells represent separate populations with distinct distributions, and thus PmCRH and its binding protein are synthesized and released by two different cell systems. The analysis of the distribution of both PmCRH and PmCRHBP at different life stages (prolarval, larval, juvenile, and adult) revealed different timings in the appearance of the different juvenile/adult populations. These developmental changes might be related to the complex/long-life cycle of lampreys: 5 to 7 years for larval development and with big changes in life after metamorphosis, from filter-feeding larvae that live burrowed in the river sediment to adult parasitic animals.

Since PmCRH and PmCRHBP are expressed by different populations, future studies should investigate how these two systems are integrated and regulated. Our results provide an anatomical basis for future functional studies on the role of CRH/CRHBP in the CNS of lampreys.

## SUPPLEMENTARY TABLE LEGEND

**Supplementary Table 1.** Animals used in each of the experiments. In some cases, serial sections from the same animal (color coded) were used for different experiments.

## FIGURE LEGENDS

**Supplementary Figure 1.** Photomicrographs of transverse sections of the spinal cord (only 1 half of the cord is shown) of adult (a-b) and larval (c-d) sea lampreys at different spinal cord levels (5^th^ gill: a and c; caudal spinal cord: b and d). Note that the dorsal column shows almost no PmCRH-ir fibers and most of them are located in the lateral portion of the spinal cord (arrows). The central canal (cc) is always to the left. Scale bars, 150 µm (a), 75 µm (b, c, d).

## Abbreviations used in Figures

ahr: anterior hypothalamic recess
cc: central canal
ch: optic chiasm
chor: choroid plexus
DC: dorsal column
DCN: dorsal column nucleus
DIG: dorsal isthmic grey
dIs: dorsal region of isthmus
dIsc: dorsal isthmic commissure
dV: trigeminal descending nucleus
fr: fasciculus retroflexus
gl: glomerular layer of the olfactory bulb
H: habenula
Hy: hypothalamus
III: third ventricle
igl: inner granular layer of OB
Ip: interpeduncular nucleus neuropil
Is: isthmus
IV: fourth ventricle
IX: glossopharyngeal motor nucleus
lH: left habenula
rH: right habenula
LP: lateral pallium
lt: lamina terminalis
Ma: mammillary region
Mes: mesencephalon
mlf: medial longitudinal fascicle
MP: medial pallium
mPO: magnocellular preoptic nucleus
MRRN: medial rhombencephalic reticular nucleus
mV: trigeminal motor nucleus
mv: mesencephalic ventricle
n: notochord
Nh: neurohypophysis
Nmlf: nucleus of the medial longitudinal fascicle
Npoc: nucleus of the postoptic commissure
NPt: nucleus of the posterior tubercle
OB: olfactory bulb
OLA: octavolateralis area
OT: optic tectum
P: pineal organ
Pc: posterior commissure
PEN: posterior entopeduncular nucleus
PO: medial preoptic nucleus
poc: postoptic commissure
PP: parapineal organ
PRRN: posterior rhombencephalic reticular nucleus
Pt: pretectum
Pth: prethalamus (ventral thalamus)
PTu: posterior tubercle
PV: paraventricular nucleus
Rho: rhombencephalon
SC: spinal cord
Se: septum
SH: subhippocampal lobe
SRRN: superior rhombencephalic reticular nucleus
St: striatum
Teg: mesencephalic tegmentum
Th: thalamus (dorsal thalamus)
TS: torus semicircularis
Tu: tuberal nucleus
V: ventricle
Vm: trigeminal motor nucleus
VII: facial motor nucleus
vIs: ventral region of isthmus
vTu: ventral tuberal nucleus
IX: glossopharyngeal motor nucleus
lH: left habenula
rH: right habenula
LP: lateral pallium
Lt: lamina terminalis
Ma: mammillary region
Mes: mesencephalon
Mlf: medial longitudinal fascicle
MP: medial pallium
mPO: magnocellular preoptic nucleus
MRRN: medial rhombencephalic reticular nucleus
mV: trigeminal motor nucleus
mv: mesencephalic ventricle
n: notochord
Nh: neurohypophysis
Nmlf: nucleus of the medial longitudinal fascicle
Npoc: nucleus of the postoptic commissure
NPt: nucleus of the posterior tubercle
OB: olfactory bulb
OLA: octavolateralis area
OT: optic tectum
P: pineal organ
Pc: posterior commissure
PEN: posterior entopeduncular nucleus
PO: medial preoptic nucleus
Poc: postoptic commissure
PP: parapineal organ
PRRN: posterior rhombencephalic reticular nucleus
Pt: pretectum
Pth: prethalamus (ventral thalamus)
PTu: posterior tubercle
PV: paraventricular nucleus
Rho: rhombencephalon
SC: spinal cord
Se: septum
SH: subhippocampal lobe
SRRN: superior rhombencephalic reticular nucleus
St: striatum
Teg: mesencephalic tegmentum
Th: thalamus (dorsal thalamus)
TS: torus semicircularis
Tu: tuberal nucleus
V: ventricle
Vm: trigeminal motor nucleus
VII: facial motor nucleus
vIs: ventral region of isthmus
vTu: ventral tuberal nucleus

## Data Sharing and Data Availability

Antibodies and histological data generated during this research are available from the corresponding author upon reasonable request.

## Funding

Grant PID2020-115121GB-I00 funded by MCIN/AEI/10.13039/501100011033 to A. Barreiro-Iglesias. Grant ED431C 2021/18 funded by the Xunta de Galicia. The European Molecular Biology Organization (EMBO) granted a long-term EMBO fellowship to D. Sobrido-Cameán (ALTF 62-2021).

## Acknowledgements

The authors thank the staff of Ximonde Biological Station for providing the lampreys used in this study, and the Microscopy Service (University of Santiago de Compostela) and Dr. Mercedes Rivas Cascallar for confocal microscope facilities and help.

## Notes

### Competing Interest Statement

The authors have declared no competing interest.

