## Supplementary figures and images for "Organization of the corticotropin-releasing hormone and corticotropin-releasing hormone-binding protein systems in the central nervous system of the sea lamprey *Petromyzon marinus*"

### Supplementary Figure 1

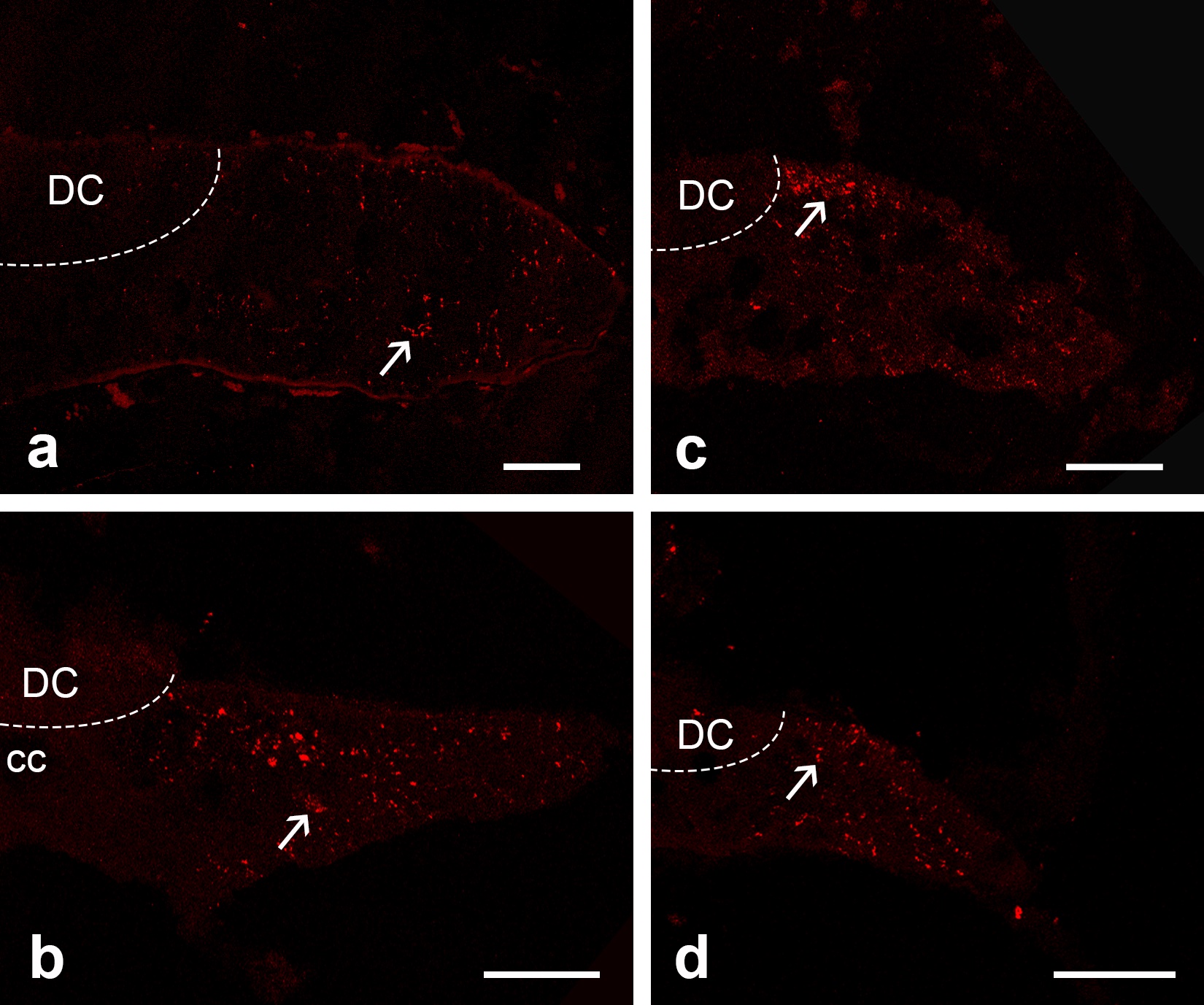
