## Supplementary Table 1 for "Organization of the corticotropin-releasing hormone and corticotropin-releasing hormone-binding protein systems in the central nervous system of the sea lamprey *Petromyzon marinus*"

| | Prolarvae | Larvae | Post-metamorphic juveniles | Upstream migrating adults |
|---|---|---|---|---|
| **Total RNA extraction** | - | 8 | - | - |
| ***PmCRH ISH*** | 15 | 3 | 4 | 3 |
| **PmCRH IHC** | 15 | 5 | 2 | 5 |
| **PmCRH + TH IHC** | - | 3 | 4 | 3 |
| **PmCRH IHC + Neurobiotin** | - | 4 | - | - |
| **PmCRH IHC + *PmCRHBP ISH*** | - | 2 | 2 | - |
| ***PmCRHBP ISH*** | 15 | 3 | 4 | 3 |
| **Total** | 15 | 20 | 6 | 8 |
